# Quantitative comparison of *in vitro and in vivo* embryogenesis at a single cell resolution

**DOI:** 10.1101/2021.05.07.443179

**Authors:** Yuqi Tan, Abby Spangler, Michael Farid, Da Peng, Patrick Cahan

## Abstract

Stem cell engineering is a powerful platform to address numerous challenges in regenerative medicine and developmental biology. Typically, engineered populations are derived by exposing pluripotent stem cells to a series of signaling cues meant to recapitulate developmental milestones, such as the induction of the primitive streak. Morphologic, genetic, transcriptomic, and functional differences between fully differentiated in vivo and engineered populations have long been recognized. However, the correspondence between engineered and native embryonic progenitors has been relatively less well characterized, largely due to their transient and highly heterogenous nature, and therefore many studies have relied on expression of a few canonical markers to ensure that their cells are on the correct path. To address this challenge, we first generated an *in vivo* gastrulation mouse single cell RNA sequencing (scRNA-seq) reference data set and used it to train a collection of computational tools for comparing cell types, states, regulators, and trajectories across studies. Then we used these tools to evaluate a commonly used protocol designed to promote mesoderm derivation, as well as four previously published biomimetic protocols. Finally, we diversified our toolkits to reach a broader scientific community by implementing our primary analysis tool in Python and as an interactive web application.

## Introduction

The embryo undergoes striking morphogenetic movements and self-assembles to form complex tissue and organs, from fertilized eggs to a gastrulating embryo. Lineage specification and axis-patterning events are guided with precision by external signals and internal expression. Through proliferation and differentiation, cells in gastrulation are differentiated into three germ layers, namely ectoderm, mesoderm, and endoderm. An array of studies has characterized morphologic, genetic, transcriptomic, and functional features for each lineage. We were interested in how engineered cells undergo gastrulation *in vitro* as compared to *in vivo*. From the past literature, we are aware that the highly heterogeneous target cell populations and frequent immature cell types are common among the directed differentiation protocols. Thus, it urges for stringent and quantitative benchmarking engineered cells against their embryonic counterparts (Bock et al., 2011; Kim et al., 2011; Osafune et al., 2008; Wu et al., 2007).

In lieu of providing new therapeutic potential, directed differentiation utilizes the intrinsic self-organizing ability of the pluripotent stem cells and allows us to study the development *in vitro*. The processes of cell differentiation, growth and movement occur simultaneously during gastrulation in a three-dimensional environment. The complexity renders it difficult to study them *in utero* in a controlled manner. Hence, a simplified system is needed to study gastrulation. Tremendous efforts have been made to generate a diversity of cell types *in vitro*. This has been done both through directed differentiation protocols, which seek to capture specific aspects of gastrulation to generate certain cell types of interest, or though more general models such as gastruloids, which aim to recapitulate the spectrum of gastrulation events in vitro, and embryoid bodies, which aims to recapitulate the early embryogenesis or gastrula *in vitro*. A recent study has also demonstrated the possibility of observing continuous embryogenesis via ex-utero culture (Aguilera). The robustness and scalability of these biomimetic studies have shown great promises in modeling human development that would not have been possible otherwise. However, how precisely these biomimetic models pass through important embryonic milestone is not yet fully understood. Furthermore, a direct comparison between these biomimetic models is absent from the current literature.

Whether we utilize directed differentiation for cellular regeneration or understanding development, a crucial question awaits to be addressed is how quantitatively similar the *in vitro* cell types to their *in vivo* counterparts. We would first require high quality *in vivo* references for direct comparisons. To circumvent the difficulties of limited materials, we proposed to generate *in vivo* mouse gastrulation at a single cell resolution. Recent addition of whole single-cell transcriptomic data on gastrulation and organogenesis further bolstered the power of our *in vivo* references (Argelaguet et al., 2019; Cao et al., 2019; Nowotschin et al., 2019; Peng et al., 2016; Pijuan-Sala et al., 2019).

With a comprehensive reference, we can conduct a number of morphologic, transcriptomic, proteomic, structural, and functional measurements to benchmark *in vitro* cell types to that of *in vivo*. Morphology is the first feature that researchers can inspect, but the comparison of such features is relatively low throughput. Functional assays, such as albumin production for hepatocytes, can assess the functionality of the engineered cells. Even though some consider functional assays as the ultimate test for the fidelity of engineered cells, it is not yet possible to devise one for every cell type. A functional assay is even more difficult to design if the engineered target is to resemble a developing embryo. While morphological and functional assays may be useful for defining and benchmarking cell identity, metrics would need to be generated independently for each cell type. Thus, a direct comparison using whole transcriptomic profiling can serve as a uniform metric across different cell types. Additionally, with advancements in computational algorithms, we can probe into the genome-wide profiles of those cells *in vitro* to connect their past, examine their current state, and infer their future trajectory using snap shots of genome-wide profiles. A number of genome-wide profiling techniques (e.g. RNA sequencing, chromatin immunoprecipitation sequencing) have been employed in the past two decades to characterize the lineage specification in multicellular organisms. However, these approaches require more starting materials than is available in one single cell, limiting their applications to cell populations. Hence, we propose to use single-cell transcriptomics to parse rare intermediates and differentiation trajectories.

Previously, we developed singleCellNet, which is a supervised machine learning algorithm to quantify the transcriptomic differences of the engineered cells and assess cell fate engineering protocols to their adult counterparts *in vivo* (Tan and Cahan, 2019). A number of single cell RNA sequencing cell typing methods, such as support vector machine (SVM) and SCMAP, have demonstrated high accuracy in annotating adult cell types (Cortes and Vapnik, 1995; Kiselev et al., 2018). Yet, none of the methods have demonstrated their accuracy in mapping embryonic cells which have more dynamic transcriptomes. Additionally, the quantitative transcriptomic comparison between engineered cell types to their developing counterparts was not possible due to the limited embryonic references available. Now that we and other independent studies have generated a few high quality scRNA-seq datasets for major cell types and lineages in mouse gastrulation and organogenesis (Argelaguet et al., 2019; Cao et al., 2019; Grosswendt et al., 2020; Ibarra-Soria et al., 2018; Nowotschin et al., 2019; Pijuan-Sala et al., 2019). Leveraging the computational tools and proper references, we can quantify the differences of their cell type identity at a single-cell resolution.

## Results and discussion

### Construction and validation of *in vivo* gastrulation classifiers

In this study, we focus our evaluations between *in vitro* and *in vivo* cells on two parts: 1) the similarity on their cell type identity and 2) the similarity on their differentiation trajectory (**Figure1A**). There are two strategies to compare the reprogramming paths of reprogrammed cells to *in vivo* cells. The first approach is to integrate *in vitro* and *in vivo* cells together with batch correction and observe how well they overlap with each other in a shared low dimensional embedding. The second approach is to employ cell typing algorithms using the *in vivo* cells as references. To examine whether we could use dimensionality reduction-based approaches to query cell types, we applied batch correction to four *in vivo* gastrulation reference data and a direct differentiation study from Spangler *et al* via Canonical Correlation Analysis (**Figure 1C** and **Supplementary Figure 2**). We observed significant overlap between the *in vivo* and the *in vitro* populations. To understand whether this overlap was reflective of the true biology, we quantified the cell identity of the *in vitro* cells using in vivo cells as references. Three of the *in vivo* gastrulation scRNA-seq datasets were integrated to construct a singleCellNet classifier (Refer to Methods for details). We then queried an independent *in vivo* study (Tan *et al*) and the direct differentiation protocol through the gastrulation classifier (Spangler *et al*). We visualized the classification score of primitive blood progenitors for both the independent *in vivo* study and the *in vitro* study (**Figure 1D**). Whether engineered cells presented with high similarity or low similarity to *in vivo* cells, the engineered cells perfectly aligned with the *in vivo* cells on the UMAP embedding space after batch correction. Thus, it is reasonable to conclude that the batch correction coerces reprogrammed cells to be overlapped with their most similar *in vivo* cells in low dimensional space, giving the illusion of reprogramming success. Thus, we proceeded with cell typing approaches to assess the fidelity of the engineered cells, instead of the batch correction approach. As for comparing the similarity of differentiation trajectories, we reconstructed trajectories for *in vivo* and *in vitro* data separately to bypass the computational complexity.

**Figure 1.**
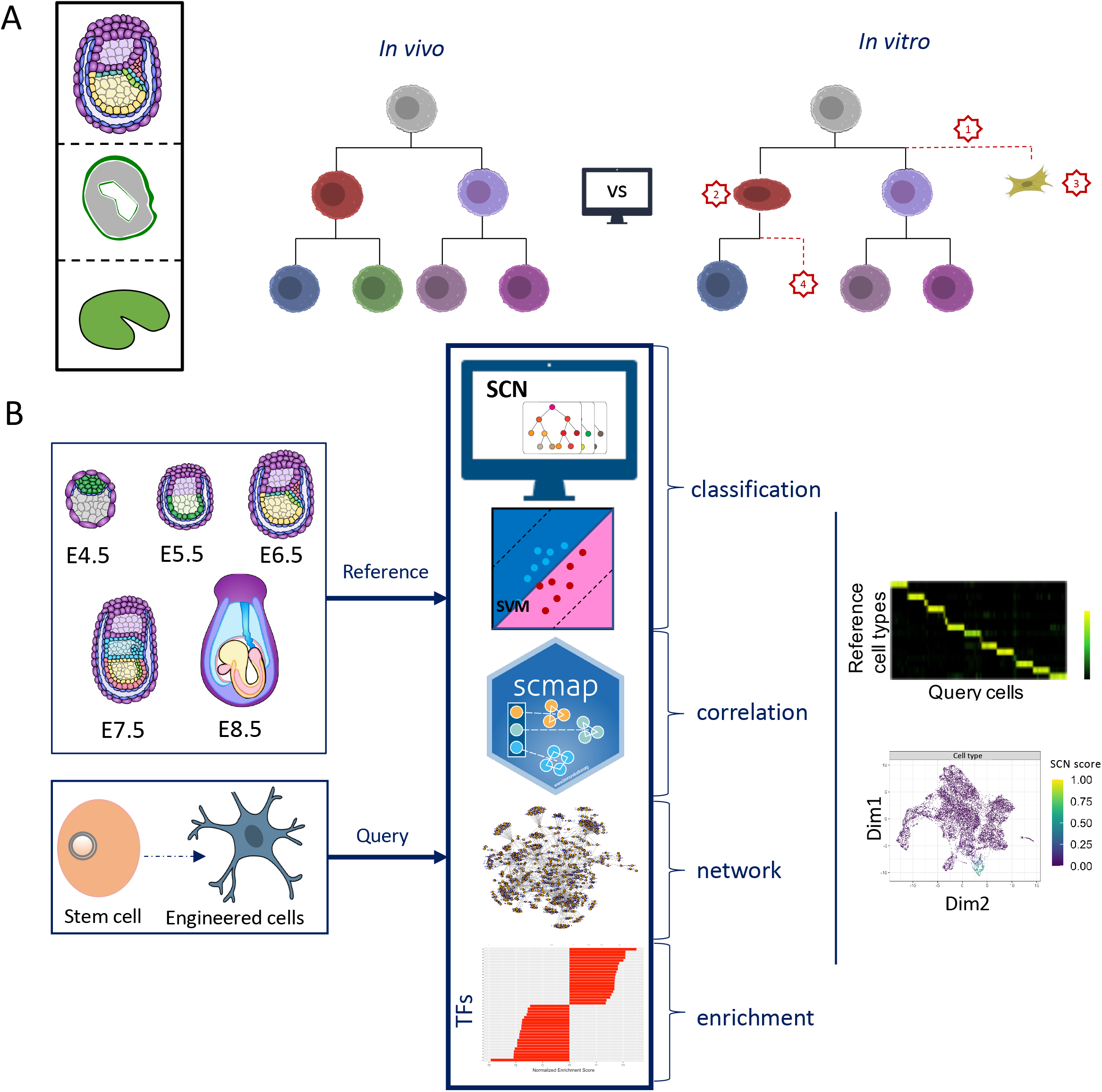
The general study design. (**A**) Our evaluations on the *in vitro* cells and *in vivo* cells will be focused on their cell identities and differentiation trajectories. (**B**) The methodological frame for the quantitative comparisons of this study. The combined meta-gastrulation scRNA-seq dataset is the reference to which the engineered cells will be compared via classification, correlation and gene regulatory network approaches. The results will be primarily visualized with heatmap and UMAP plot.

We leveraged classification, correlation, gene regulatory network (GRN), and gene set enrichment analysis (GSEA) to quantitatively evaluate cell type similarity between *in vitro* and *in vivo* cells (**Figure 1A-B**). To provide the quantitative scoring for major cell types from preimplantation to gastrulation, we integrated three different single-cell mouse gastrulation studies (Grosswendt et al., 2020; Nowotschin et al., 2019; Pijuan-Sala et al., 2019) and built a gastrulation meta-reference classifier via SCN (Tan and Cahan, 2019) (**Table1**). A total of 102,908 cells of 47 distinct cell types were integrated from the three studies to build the classifiers, namely SCN, SVM, and SCMAP-cluster meta-gastrulation classifier. We trained the SCN meta-gastrulation classifier and optimized the parameters to obtain the best performing classifier (**Supplementary Figure 3A-C**). We have selected the best performing SCN meta-gastrulation classifier which achieves 0.91 mean Area Under Precision-Recall Curve (AUPRC, a measure of specificity and sensitivity) and 0.87 Cohen’s Kappa (a chanced corrected accuracy). We used the default parameters for SVM and SCMAP-cluster.

**Table 1.** Gastrulation Reference Datasets.

| Dataset | Cell types/stages | Seq | Number of cells |
| --- | --- | --- | --- |
| Grosswendt | Epiblast, Definitive endoderm, Ectoderm early 1, Trophoblast stem cell, Parietal endoderm, Ectoderm early 2, Visceral endoderm early, Anterior primitive streak, Distal extraembryonic ectoderm, Differentiated trophoblasts, Primitive streak early, Hematopoietic/endothelial progenitors, Primitive blood progenitors, Visceral endoderm late, Neural crest, Primordial germ cells, Allantois. Posterior lateral plate mesoderm, Node, Gut endoderm, Primitive steak late, Preplacodal ectoderm, Neural ectoderm posterior, Anterior paraxial mesoderm, Surface ectoderm, Amnion mesoderm early, Neural ectoderm anterior, Secondary heart field, neuromesoderm progenitor early, Somites, Primitive blood early, Notochord, Primitive heart tube, Angioblasts, Presomitic mesoderm Amnion mesoderm late, Similar to neural crest Primitive blood late, Neuromesodermal progenitor late, Fore/midbrain, Future spinal cord, Pharyngeal arch mesoderm | 10x | 88779 |
| Notowschin | Primitive endoderm<br>Epiblast<br>Trophectoderm<br>Inner cell mass<br>Trophoblast stem cell<br>Definitive endoderm<br>Visceral endoderm early | 10x | 12556 |
| Pijuan-Sala | Extraembryonic endoderm<br>Extraembryonic mesoderm | 10x | 1573 |

To evaluate and calibrate the meta-gastrulation classifiers, we sampled three gastrulation time points from wild type C57BL/6J mouse gastrula (E6.5, E7.5, E8.25) (**Figure 2A-C and Supplementary Figure 4**). A total of 11,726 single-cell transcriptomes were profiled via 10x Genomics with a median of 5,514 genes per cell and a median of 36,721 UMI counts per cell. After removing low-quality cells and performing standard preprocessing using the standard SCANPY pipeline (Wolf et al., 2018), we retained 11,102 single-cell profiles for further analyses and identified 20 clusters. To unbiasedly assess the performance of the classifier, we annotated the clusters using cell type specific markers and captured 16 distinct cell types (**Supplementary Notes 1**). To show that our annotation was in accordance with the literature, we overlaid a few cell-type specific marker gene expressions with our clustered cells on a UMAP embedding (**Figure 2C**). FgF5, a widely used epiblast marker was expressed primarily in cluster 8. Nkx1-2, a known neural mesodermal progenitor (NMP) marker, was also primarily expressed in cluster 3 and cluster 13. T, a marker typically associated with notochord development, was strongly expressed in cluster 12,1. Similarly, Lama1, a parietal endoderm (ParE) marker also showed high specificity to cluster 19.

**Figure 2.**
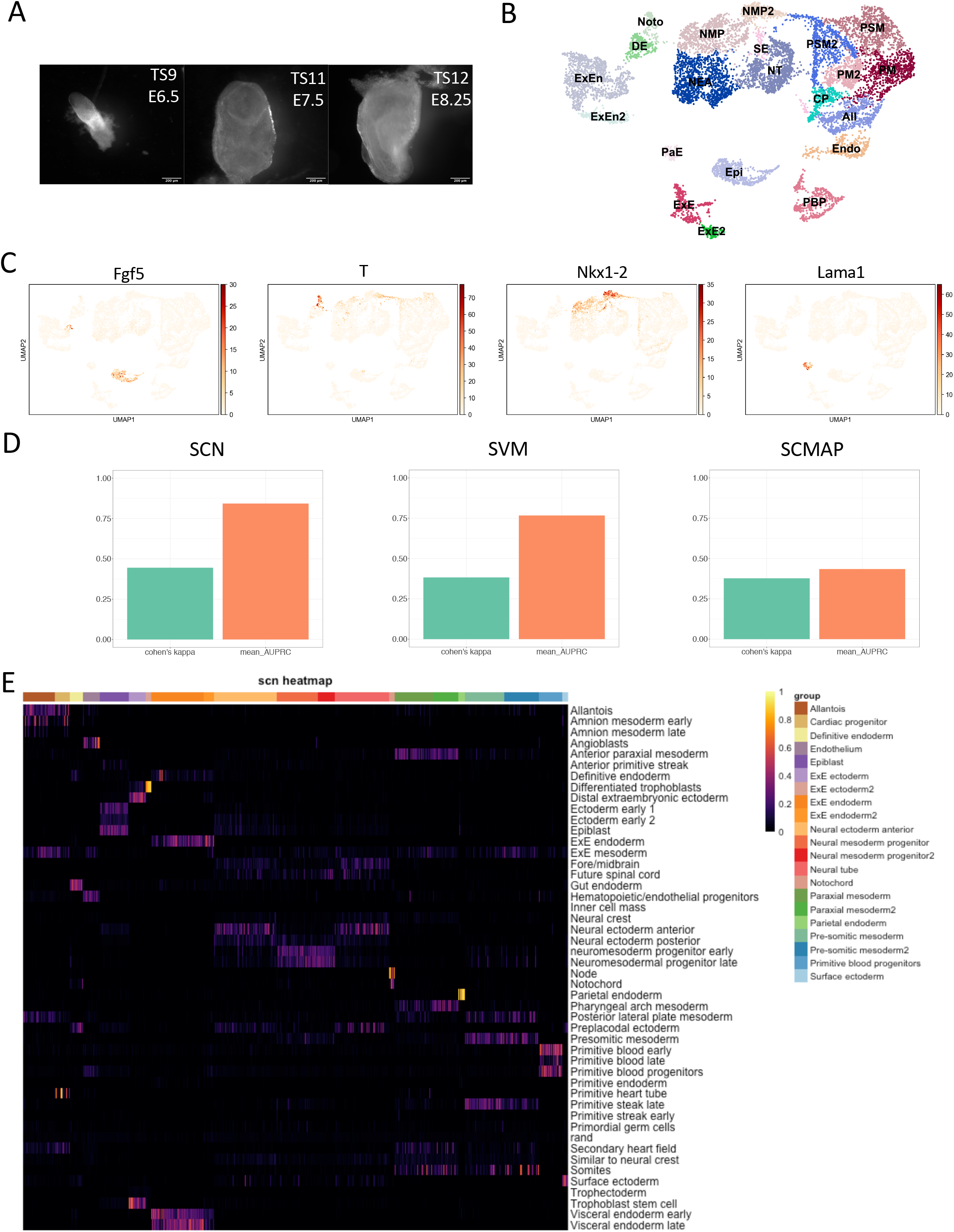
A single-cell resolution of mouse gastrulation. (**A**) Representative images of gastrulating embryos and their stages (scale bar 200μm) (**B**) A UMAP embeddings of all 11,102 scRNA-seq profiles. Clusters were annotated by cell-type specific marker gene expression. (**C**) Selective cell type-specific marker gene expression overlaid on the UMAP embeddings. (**D**) Assessments of the performance of meta-gastrulation classifiers constructed with SCN, SVM and SCMAP are shown in the barplots. Two metrics, Cohen’s kappa and Mean Area Under Precision Recall Curves, are measured. (**E**) The full SCN classification heatmap for the in-house gastrulation dataset is generated with the meta-gastrulation SCN classifier. The x-axis displays the SCN classifier categories, and the y-axis are single cell profiles that are clustered by their marker-gene based annotation. The SCN classification score ranges from 0 to 1. The higher the score the higher the transcriptomic similarity to the corresponding classifier category.

Using this marker gene-based annotation as ground truth, we applied the optimized SCN, SVM, and SCMAP-cluster meta-gastrulation classifier to the annotated single-cell gastrulation data (**Figure 2D**). Since none of the methods have applied to embryonic cell typing before, thus we used our dataset as an external validation dataset to examine how well these tools perform in handling embryonic cell typing (**Supplementary Figure 4G-I**). We were only able to assess 12 out of 16 cell types that have the matching reference cell types in the meta-gastrulation classifiers. The best classification performance using our embryo data for external validation was SCN, closely followed by SVM and then SCAMP-cluster. 11 out of 12 classifier categories that we were able to assess, achieved satisfactory performance (AUPRC >0.5) for both SCN and SVM, while SCMAP shows lower sensitivity. We also experimented with integrating different methods using a weighted average scoring approach, but the naïve weighting approach did not improve the classification performance (**Supplementary Figure 5**). Hence, we decided to rely on SCN classification as our primary assessment tool in the following analyses and supplemented with SVM and SCMAP. As we looked into the reason for the lower performance in the definitive endoderm (DE) classifier category, we noticed that most of the DE cells were being classified as gut endoderm (GE), which is a derivative of DE (**Figure 2E**). The notochord cluster contained node cells as well, which are the progenitors of the notochord. Collectively, the result shows that 1) SCN meta-gastrulation classifier performs well and 2) the classification-based annotation was more sensitive in distinguishing closely related cell types than cluster-based annotation using cell-type-specific markers. We later used this optimized classifier to assess ten cell fate engineering studies (**Table 2**) and four biomimetic studies (**Table 3**).

**Table 2.** Biomimetic Model Datasets.

| Dataset | Major cell types | treatment | Seq | # cells | Time points | Study |
| --- | --- | --- | --- | --- | --- | --- |
| EB_multiseq | All, cardiac progenitor, endothelium , EPI, ExE, ExEn Endo, NMP, NE, PM, PsM, SE, blood progenitor (BP) | Wnt3a and Activin A along with : Gsk inhibitor, or BMP4 for primitive streak induction at day2, bFGF at day4, bFGF and Gdf5 at day6 | 10x | 12335 | D4, D5, D6, D7, D8 | This paper |
| EB_recording | PS, PrE, EPI, BP, Endo, ESC, PGC, Mes | mESC media without LIF with feeder MEFs | CEL-seq2 | 3489 | D0, D2, D4, D6, D8, D10, D12, D14 | Kim et al., 2020 |
| EB_replicate | BP, Endo, Ect, PGC, Mes | mESC media without LIF with feeder MEFs -30min tamoxifen at day8-9 to induce CreERT2 | CEL-seq2 | 1281 | D14 | Kim et al., 2020 |
| gastruloid | Smt, cardiac, PM, differentiated smt, PsM, NMP, mesenchyme , endothelium , All, PGC, Endo | N2B27 medium, 3 μM chiron pulse at day2, embedded the cells in Matrigel at day3 | 10x | 25202 | D5 | Van der Brink et al, 2020 |

### Combinatorial morphogen induction of mesoderm

Our next step was to test the utility of the meta-gastrulation classifier to investigate mesoderm formation *in vitro*. Previously, we induced mESC differentiation with morphogens *in vitro* to form EBs and to induce primitive streak formation (Craft et al., 2013; Spangler et al., 2018). subjecting this data to SCN analysis revealed that many of the derived cells at day 4 and day 6 exhibited a neuronal signatures, and one cluster had a high similarity to the notochord (**Supplementary figure 5**). In this study, we followed a similar protocol for primitive streak (PS) induction, but tested how modulating BMP and Wnt signaling would impact germ layer derivatives. We tested induction with WAG (Wnt3a, Activin A, Gsk inhibitor) and WAB (Wnt3a, Activin A, BMP4) on day 2 for 48 hours, followed by bFGF on day 4 (**Figure 3A**). Since Brachyury marks PS, we also sorted based on Bry-GFP on day 4, and reaggregated the embryoid bodies for later sequencing. The control groups remained unsorted. To examine whether the modified protocols increased the yield of mesodermal lineage, we applied scRNA-seq to cells in day 4 through day 8 (five time points) and obtained 12,335 single-cell transcriptomes with a median UMI count of 12,481 (**Supplementary Figure 6A**). After stringent preprocessing, we identified 18 distinct clusters. However, because of the stress FACS imposed on cells, we recovered fewer cells in the sorted samples compared to the unsorted ones (**Supplementary Figure 6B**).

**Figure 3.**
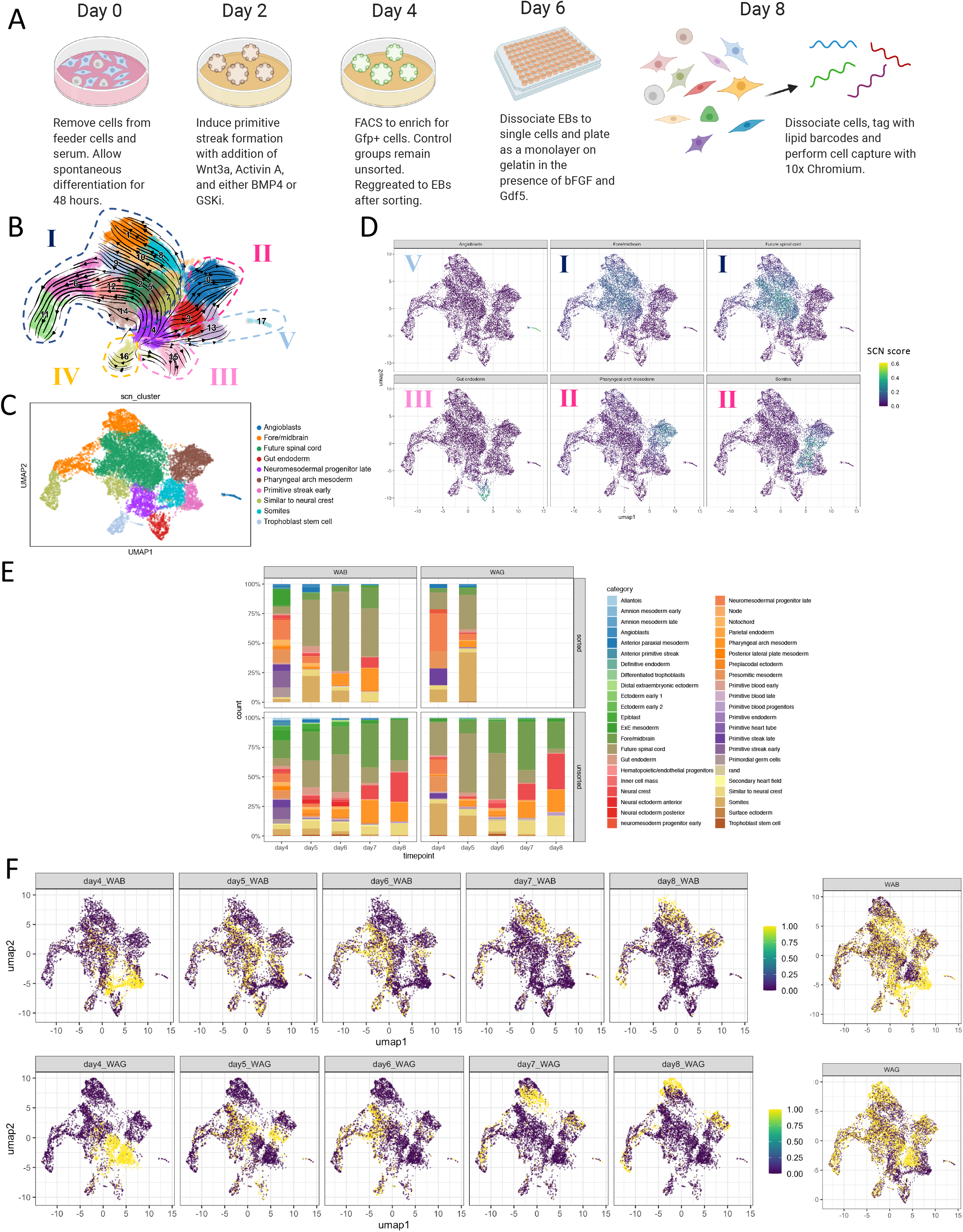
A time-course experiment of an embryoid-body based directed differentiation study at a single-cell resolution. (**A**) The experiment design for the culture and treatment of differentiating mESCs towards mesoderm cell types through forming embryoid bodies. Two primitive streak induction treatments were employed. (See Methods for details). scRNA-seq profiles were sampled from day 4 to day 8 at a 24hr interval. (**B**) A scVelo plot of potential differentiating trajectories of the differentiating mESCs. The arrows are velocities derived from the dynamical model for mESCs differentiation. (**C**) A UMAP embeddings of all EB scRNAseq. Clusters are annotated by the most cell types classified by SCN in that cluster. (**D**) UMAP embeddings with SCN colored with SCN classification score from the meta-gastrulation classifier. The scores of six cell types are shown here. SCN classification score ranges from 0 to 1. (**E**) An attribution plot showcases the cell type compositions between sorted and unsorted groups over the course of EB differentiation and under two primitive streak induction treatment. (**F**) Cells are first stratified by timepoints and treatment and then are shown on the UMAP embeddings. Only the unsorted cells for each condition were plotted. (bottom-right) A agglomerate of UMAP embeddings of cells per treatment.

To examine how each cell relates to each other, we used scVelo to infer the trajectories of all 12,335 scRNA-seq profiles (Bergen et al., 2020). We identified cluster 4 as the root of the trajectory and there were five major fates (**Figure 3B**). We queried the embryoid body (EB) cells through the SCN meta-gastrulation classifier. Combining the inferred trajectories with the SCN classification result (**Figure 3C**), we concluded the following observations (**Figure 3D**). The majority of the cells acquired the neural fate (Fore/midbrain and future spinal cord) (Fate I). Second largest cell group was differentiated towards pharyngeal arch mesoderm (PhM) and somite (Fate II). A portion of the EB cells adopted GE fate (Fate III). A few EB cells differentiated towards the angioblast fate (Fate V). Meanwhile, we saw the emergence of a group of stagnant reprogrammed cells, where most of them were classified as inner cell mass (ICM), epiblast (EPI), and trophoblast stem cells (TSC) (Fate IV). (**Supplementary Figure 6C**).

To investigate how the WAB and WAG treatment influence lineage propensity, we analyzed the temporal fate transition of the EB cells. We observed that sorting by bry-GFP+ does not enriched for mesodermal populations (**Figure 3E**). The additional sorting also induced great stress to EB cells and lowered their viability (**Supplementary Figure 6D-H**). Since there was much variability of cell number after being sorted, the treatment comparison was conducted between the unsorted groups. We observed that the WAB treatment yielded more angioblast, GE, and future spinal cord (FSC) cells (**Figure 3F**). Meanwhile, WAG induction produced more differentiated Fore/midbrain (FM) cells and somite. On day 4, both WAG and WAB treatments successfully induced PS cells, while most somitogenesis was observed only under WAG treatment. On day 5, the formation of angioblasts peaked in both treatments. On day 6, with a small fraction of the cells becoming GE and PhM, the majority of the EB cells were fated to become FSC. From day 6 to day 8, while the majority of the cells in both treatments were specified to become FM, we can also observe the steady development of more differentiated PhM.

We noticed that WAB treatment encouraged more EB cells to acquire GE and angioblast fate, while WAG treatment facilitates EB cells to undergo somitogenesis. The majority of the directed differentiation cells still adopted the ‘default’ neuroectodermal cell fate. To enrich for mesodermal cell type, a negative enrichment against neural cell markers could be implemented in the future. As compared to their *in vivo* counterparts, the engineered cells exhibited more prominent hybrid identities and therefore there is much room for improvement in precision for the directed differentiation approach.

### Quantitative evaluation of biomimetic studies to gastrula

After classifying our direct differentiation study with the meta-gastrulation classifier and comparing the *in vitro* and *in vivo* trajectories, we wanted to expand our comparisons to publicly available biomimetic studies. We have selected four scRNA-seq biomimetic studies (three EB-based and one gastruloid-based) to evaluate the differences between the two approaches in modeling gastrula (**Table 2**). To align *in vivo* stages to the *in vitro* timepoints between the two approaches, we created the schematic based on the original reports (**Figure 4A**) (Beccari et al., 2018; van den Brink et al., 2020; Kim et al., 2020). For easier recognition, we named the four biomimetic studies as follows: EB_multiseq, EB_recording, EB_replicate, and Gastruloid. EB_multiseq was generated in this study as described in Figure 3. The other studies were obtained from published papers. The details on how these protocols were generated can be found on Supplementary note 2.

**Figure 4.**
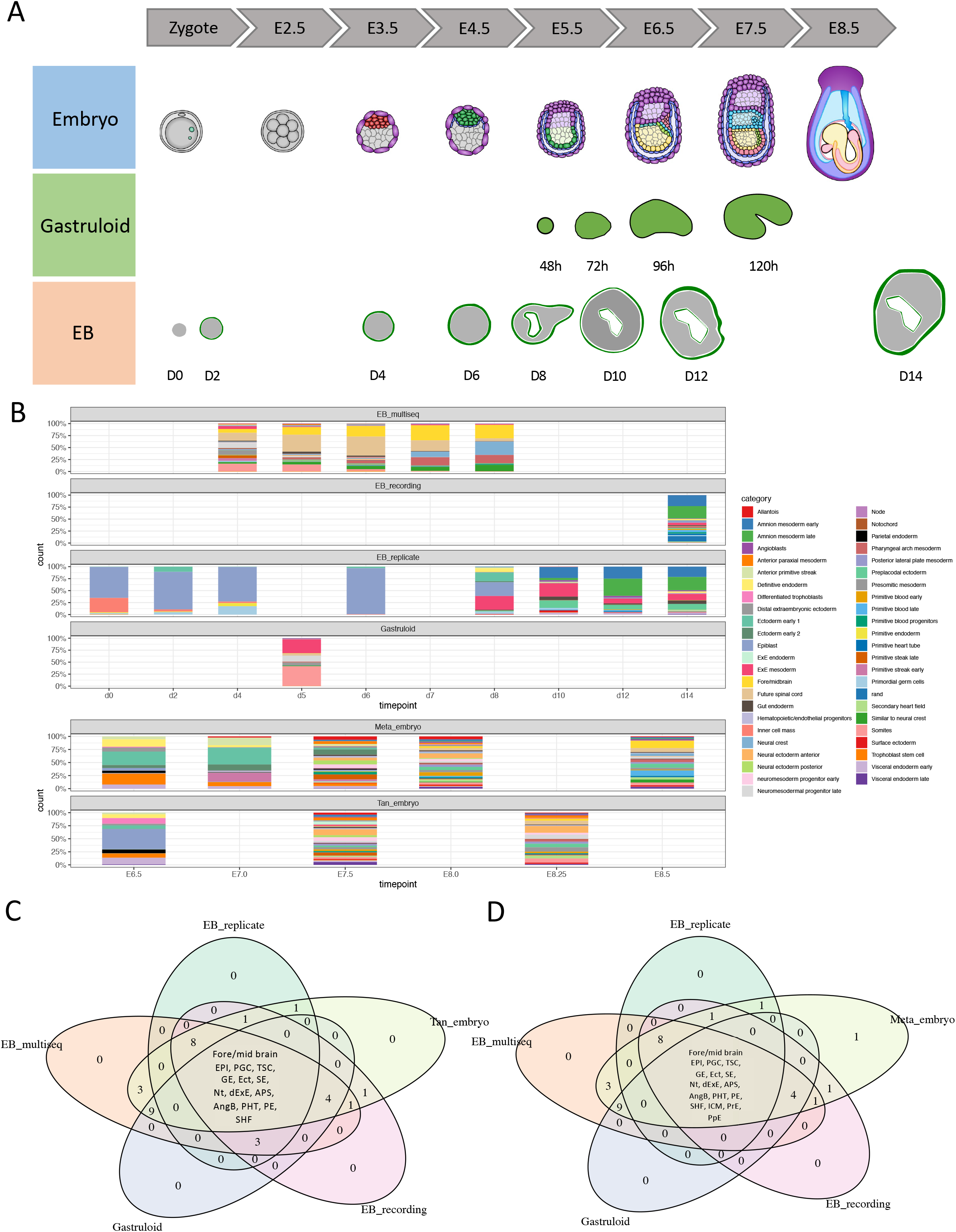
Cross-study comparison of biomimetic studies. (**A**) A literature-based schematic to align i*n vitro* timepoints for each biomimetic approach and their corresponding *in vivo* timepoints (**B**) An attribution plot of four biomimetic studies colored by their annotation from the SCN meta-gastrulation classifier. The attribution is stratified by timepoints and studies. (**C**-**D**) Venn diagrams are shown here to understand the overlap between *in vitro* and *in vivo* cell type composition.

To quantify how cellular composition differs between biomimetic approaches, we applied the SCN meta-gastrulation classifier to four biomimetic studies and visualized the cell type composition for each study (**Figure 4B**). We observed two major trends from the SCN cell-type proportion analysis.

The first observation is that even within the same type of engineering approach there are high variability in lineage propensity. The EB_multiseq study has shown a high percentage of neuro-epithelialization (FM, FSC, and NC cells). The cells from EB_replicate and EB_recording demonstrated strong propensity towards becoming amnion mesoderm. In EB_replicate, we saw an increase in EPIs which peaked on day 6. EB_replicate began to diversify in cell composition from day 8. Even though we have observed all three germ layers on day 8, the extraembryonic lineage, particularly ExM and amnion mesoderm, expand proportionally faster than other lineages. Similar cell type composition was found on day 14 for the EB_recording study. A significant percentage of gastruloid on day 5 is ExM. Consistent with the authors’ report, an even greater percentage of gastruloid cells on day 5 is fated to be somite. The third and fourth largest portion of cells is fated to become the Fore/midbrain and future spinal cord, which is similar to that of EB_multiseq.

The second observation is that the *in vitro* biomimetic studies show less cell type diversity than that of the *in vivo* gastrulating embryos. Fewer than twenty out of forty-seven cell types profiled from the gastrulating embryos were generated *in vitro* (**Figure 4C-D**). The only cell type that was not reproduced in the *in vitro* system was trophectoderm. To align the *in vitro* time point to *in vivo* embryonic day, we computed the similarity with the cell type composition via cosine similarity (**Supplementary Figure 7**). From this analysis, we observed that the literature proposed time point alignment between the biomimetic studies and *in vivo* embryo is not entirely accurate. We compared twelve embryonic time points from Grosswendt *et al* gastrulation data to the *in vitro* studies (Grosswendt et al., 2020). The cell type compositions of all five time points from the EB_multiseq study resembled highly to that of the E8.0 gastrula. The similarity trend decreased overtime. Day 14 sample from EB_recording shared the highest similarity of the cell type composition to that of the E8.0 gastrula. Meanwhile, day 0 to day 8 timepoints from the EB_replicate study better aligned to E5.5. From day 8 to day 10, we noticed the cell type compositions started to resemble more towards E7.75 and E8.25 gastrula. The resemblance continued but with decreased correlation overtime. The cell type composition of day 5 gastruloid shared the resemblance to that of E7.75 and E8.25 gastrula.

To further investigate on what are the differences in range of classification scores between *in vitro* and *in vivo* cells of a given cell type, we visualized the classification score in a density plot to showcase its continuous nature (**Supplementary Figure 8**). We further included the held-out training data and our gastrula scRNA-seq data as references and performed Wilcoxon Rank Sum test. In the statistical analysis, we used both the held-out data and our gastrula scRNA-seq data as references respectively. As expected, none of the methods has generated engineered cells that are indistinguishable from that of the held-out data. However, we noticed both EB and gastruloids approaches have generated a few cell types, such as angioblasts, that are statistically indistinguishable from that of our independent gastrula reference data (**Figure 5A-B**).

**Figure 5.**
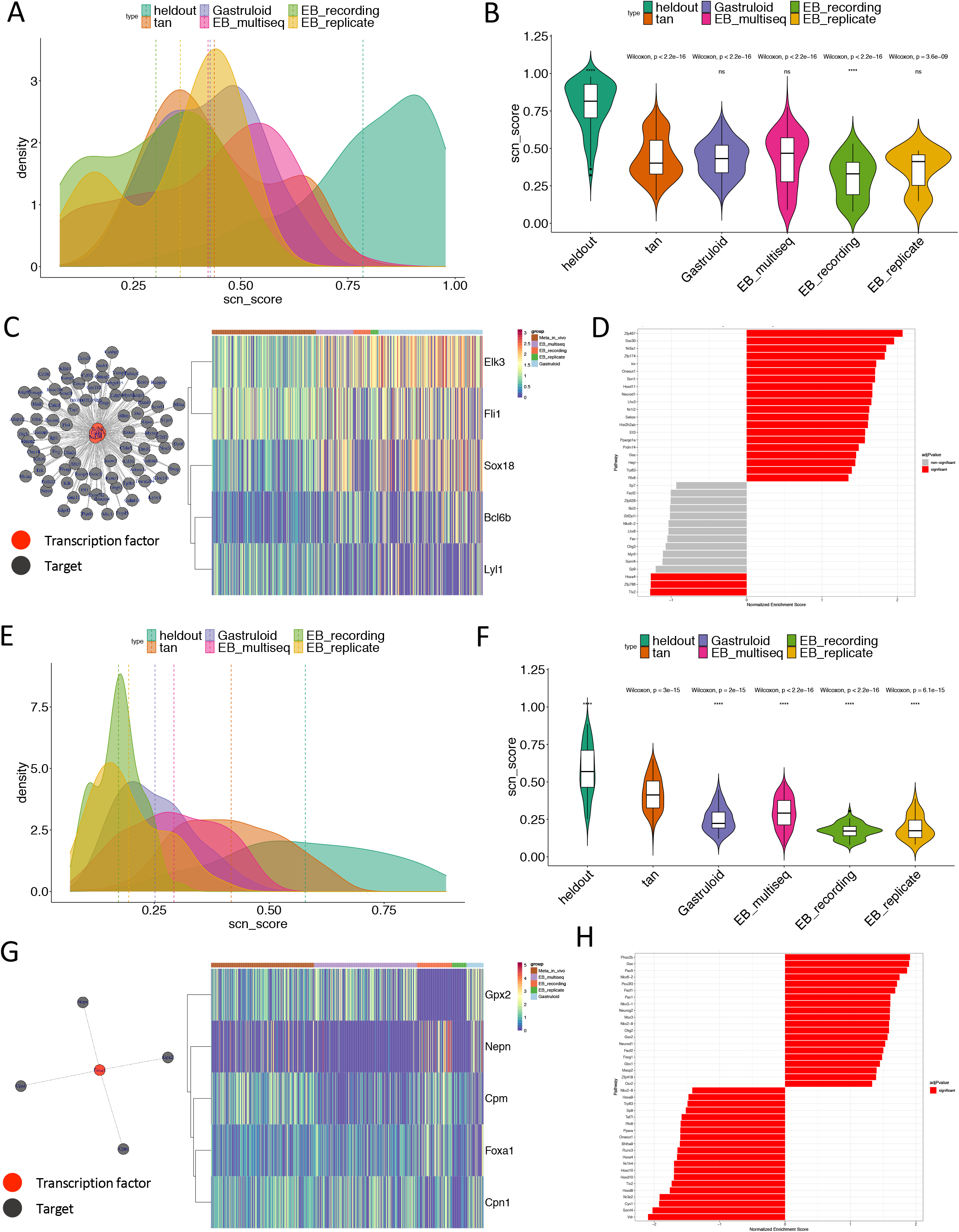
Compare biomimetic cells to gastrulating cells using single-cell network enrichment. (**A**) A SCN classification distribution plot showcases the classification score for angioblasts by the SCN meta-gastrulation classifier across four biomimetic studies (**B**) The same classification scores are plotted in violin plots. The differences amongst the distributions of the classification scores are tested with Wilcoxon’s Rank Sum test. The numeric number of p-values denotes the comparison when the heldout distribution was the control for the comparison. The asterisks or ns denotes test results where Tan embryo dataset, an independent gastrulation scRNA-seq dataset, is used as the reference. When P < 0.05, we considered there is no statistical difference between the query and the control. (**C**) Angioblast specific gene regulatory network is extracted from single-cell gastrulation GRN. The expression levels of key transcription factors for *in vivo* and *in vitro* angioblasts are shown in the heatmap. (**D**) The network enrichment analysis was performed on the list of transcription factor subnetworks that were identified from the single-cell gastrulation GRN. Significantly enriched transcription factor networks are shown in red. (**E**) A SCN classification distribution plot showcases the classification score for gut endoderm by the SCN meta-gastrulation classifier across four biomimetic studies. (**F**) The same classification scores are plotted in violin plots. The differences amongst the distributions of the classification scores are tested with Wilcoxon’s Rank Sum test. (**G**) Gut endoderm specific gene regulatory network is extracted from single-cell gastrulation GRN. The expression levels of the key transcription factor and its targets for *in vivo* and *in vitro* angioblasts are shown in the heatmap. (**H**) The network enrichment analysis was performed on the list of transcription factor subnetworks that were identified from the single-cell gastrulation GRN. Significantly enriched transcription factor networks are shown in red.

In addition to classification assessment, we employed single-cell network enrichment to further investigate the molecular similarity or dissimilarity between the *in vitro* and *in vivo* cells. We first constructed a single-cell GRN from Grosswendt *et al* gastrulation data with a Context-likelihood of Relatedness (CLR) algorithm (Faith et al., 2007). A network of five transcription factors (Bcl6b, Elk3, Fli1, Sox18, Lyl1) was extracted to be the angioblast specific GRN (**Figure 5C**). Both Sox18 and Lyl1 have previously been reported to play an essential role in angiogenesis and blood vessel maturation (Darby et al., 2001; Deleuze et al., 2012; Pirot et al., 2010). We examined the expression of this network of these five transcription factors in the four biomimetic studies and the *in vivo* reference and found the patterns of the transcription factor network are coarsely similarly expressed among the studies. One noticeable downregulation of Sox18 was observed in EB_recording, which could be the reason why angioblasts from this study had statistically lower SCN classification scores. To broadly investigate the potential dysregulated transcription factors in the EB_recording study, we identified the both positively and negatively enriched transcription factors that were identified from our gastrulation GRNs (**Figure 5D**). Top 20 and bottom 20 most differentially enriched transcription factors in the EB_recording angioblasts were shown as bar plots. Amongst the top positively enriched transcription factors, *Sox30*, and *Nr5a1* are involved in embryonic germ cell differentiation (Bashamboo et al., 2016; Zhang et al., 2018). This observation is consistent with authors’ application of their EB protocol to investigate the PGC-EPI decision. One possible explanation is that the external stimuli that favors PCG formation can extend its influences into gene expression of other cell types. Other positively enriched transcription factors are also involved in liver gene expression (*Onecut1* and *Scrt1*), forelimb morphogenesis (*Hoxd11*), and neuronal differentiation (*Neurod1* and *Lhx3*). Meanwhile, the top negatively enriched transcription factors are associated with anterior-posterior pattern (*Hoxa4* and *Sp9*). These transcription factors can serve as potential candidates for further maturing the EBs to resemble gastrula by enhancing anterior-posterior patterning. To better interpret the molecular difference biologically between the EB_recording angioblasts and the in vivo angioblasts, we performed gene set enrichment analysis based on the differentially expressed genes. We noticed that the *in vitro* angioblasts exhibited enhanced mobility (**Supplementary Figure 9A-B**).

While a few cell types, such as angioblasts, have statistically undistinguishable classification scores when compared to their *in vivo* counterparts, most *in vitro* cell types have significantly lower classification scores than that of *in vivo*. We analyzed the *in vitro* GE as an example (**Figure 5E-F**). All four biomimetic studies showed significantly lower score than the held-out data and the independent Tan embryo data. The network of a single transcription factor, *Foxa1*, was extracted to be the GE-specific GRN from the single-cell gastrulation GRN (**Figure 5G**). Gastruloid study showed noticeable downregulated *Foxa1* expression. While the *Foxa1* expression level was comparable across the *in vitro* and the remainder of *in vivo* studies, *Nepn* and *Gpx2* expression varied greatly among the *in vitro* cells. To broadly investigate the potential dysregulated transcription factors in the EB_multiseq study, we performed single-cell network enrichment using the same gastrulation GRNs as mentioned above. The top positively enriched transcription factors for EB_multiseq are predominantly related to neurogenesis (*Phox2b*, *Gsc*, *Nkx6-2*, *Pou3f3*, *Gsx2*, *Olig2*, *Neurod1*). The trend that the imprints of transcription factors that lead the specification for majority cells *in vitro* can be found in other *in vitro* cell types where they normally are not expressed holds true for rest of the biomimetic studies (**Supplementary Figure 10**). The gene set enrichment analysis based on the differentially expressed genes between EB_multiseq and *in vivo* GEs also shows positively enriched pathways related to neuronal development (**Supplementary Figure 9C-D**).

In summary, through analyzing the biomimetic studies, we observed three trends. The first observation is that the *in vitro* biomimetic studies show high heterogeneity in lineage propensity. The second observation is that the *in vitro* biomimetic studies show fewer cell type diversity than that of the *in vivo* gastrulating embryos. The third observation is that the imprints of transcription factors that leads the specification for majority cells *in vitro* can be found in other *in vitro* cell types where they normally are not expressed holds true for rest of the biomimetic studies.

### Broaden the accessibility of SCN via python implementation and an interactive web application

To expand the reach of the tool to a wider community, we implemented SCN in python (**Figure 6A**). We have shown a ten times reduction in training time with the python implementation particularly when the training dataset is large and more complex (**Figure 6B**). Additionally, we have designed a cloud-computing supported, user friendly and interactive web application for SCN where users can upload their data and visualize their SCN results interactively with the web.

**Figure 6.**
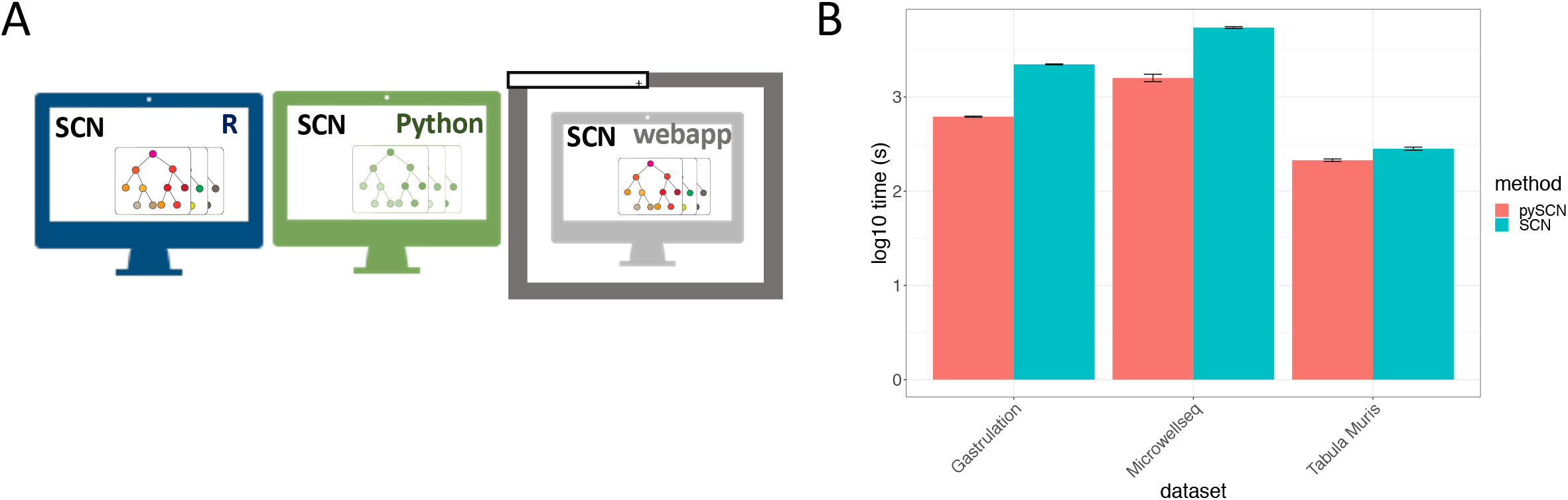
New singleCellNet implementations in Python and Web application. (**A**) Schematic for available SCN platforms (**B**) A box plot demonstrates the training time difference for three mouse atlas datasets in log10(s) scale for SCN and pySCN. Each training was bootstrapped for 10 times.

## Conclusion

In this study, we used scRNA-seq to characterize the similarity of embryonic biomimetic cell populations to their native counterparts. To do so, we incorporated three scRNA-seq gastrulation studies to form a meta-gastrulation reference. Leveraging this reference, we quantified the transcriptomic differences using classification, correlation, gene regulatory network and gene set enrichment analysis. To calibrate the performances of all the computational analysis, we have generated an independent gastrulation dataset with 11,726 scRNA-seq profiles. This new scRNA-seq gastrulation atlas will contribute to the field as an additional valuable resource to understand mouse gastrulation. Finally, from our analysis on the biomimetic studies, we have identified high variability in lineage propensity between protocols. Compared to the *in vivo* gastrulation cell types, only few cell types *in vitro* show statistically similarity in transcriptome which is quantified by SCN classification score. We also identified a gene regulatory network of five transcription factors that define the regulatory network of angioblasts.

Limited by the availability of references and high throughput data, our analyses solely focus on the comparisons of whole transcriptomes. As aforementioned, there are multiple aspects defining a cell type. The evaluations of those aspects will also be crucial to complete comprehensive assessments. The second limitation of our study is that our *in vivo* reference is annotated based on prior literature, which could prevent us from recognizing novel lineage specifications.

Despite these limitations, we believe our meta-analyses took a step forward in evaluating cell fate engineering studies and biomimetic studies with multiple lineages by direct comparisons to endogenous development. We hope to use these findings and analytical approaches to provide others with powerful insights into the assessment and improvement of future cell fate engineering experiments.

## Methods

### ESC maintenance and differentiation

Braychyury-GFP mESC cells (Gadue et al., 2006) were maintained and differentiated through day 2 of the directed differentiation protocol according to Spangler *et al*. (Spangler et al., 2018). On day 2, four different primitive-streak induction treatments were established by adding growth factors Wnt3a (25ng/mL) and Activin A (WA; 9ng/mL) along with one of the following additional growth factors: Gsk inhibitor (10mM), or BMP4 (0.5ng/mL). After a 48-hour primitive streak induction, embryoid bodies (EBs) were dissociated to a single cell suspension by incubating them in TrypLE and straining through a 40uM cell strainer. The GFP+ population was isolated by fluorescence-activated cell sorting (FACS) and subsequently re-aggregated to form EBs in SFD with bFGF (10ng/mL) for 48 hours. On day 6, EBs were dissociated and plated as a monolayer on 0.1% gelatin-coated tissue culture plates in SFD with bFGF (10ng/mL) and Gdf5 (30ng/mL) for 48 hours. Separate experiments were started in a staggered manner such that on the day of sequencing we had samples representing every day of the experiment from day 4 to day 8. We had samples representing four different primitive streak inductions conditions as well as samples that were not sorted for GFP on day 4. On the day of sequencing, EBs and monolayers were dissociated to a single cell suspension through the use of TrypLE and 40uM cell strainers.

### Single cell RNA sequencing of ESC differentiation and mouse gastrulation

For the ESC differentiation experiment, we utilized the MULTI-seq sample barcoding and library preparation protocol from the McGinnis lab (McGinnis et al., 2019) to sequence 27 samples in two 10x capture runs. In short, the protocol involved tagging cell membranes with sample-specific barcodes using a lipid-modified oligonucleotide (LMO). The LMOs (reagents obtained from the McGinnis lab) anchor into the cell membrane and allow for the attachment of a normal ssDNA oligonucleotide (MULTI-Seq barcode). A unique MULTI-seq barcode was added to each sample after which, all samples were pooled and sequenced as if they were one sample. An equal number of cells from each sample were combined to make up the pooled samples and ensure equal representation of each sample after down-stream sequencing. MULTI-Seq barcodes were used down-stream to identify which cells were from which samples. After cells were successfully tagged and pooled, they were submitted to the sequencing core facility for 10x capture and library preparation. A Truseq library preparation was performed with the necessary adjustments made to accommodate the MULTI-Seq platform. Libraries were sequenced on Illumina NovaSeq6000.

C57BL/6 wild type embryos of gastrulation stages (E6.5, E7.5, and E8.25) were collected and dissociated in TrpLE at room temperature for 10min. The dissociation was stopped with DMEM and 10% FBS. The cells were spanned down at 350rcp for 5min and resuspended in PBS. The cells were washed once more time with PBS. The cells were resuspended in PBB with 0.4% BSA before checking for cell viability and density. Standard 10x Sequencing protocol was then applied to the single-cell suspension with higher than 90% viability.

### Processing single cell RNA sequencing data

The scRNA-seq data for the ESC differentiation and the mouse gastrula was analyzed using the Scanpy package (v.1.6.0) (Wolf et al., 2018). For the mouse gastrula experiment, cell barcodes with more than 1,000 transcripts and fewer than 7,100 genes were selected. Genes detected in fewer than ten cells were excluded. Expression levels for each cell were normalized to 10,000 transcripts and logarithmically transformed. Highly variable genes were defined as those with a mean expression value between 0.0125 and 5, and with 0.5 dispersion, and used to generate the UMAP plots. Cells were clustered using the Leiden algorithm. For the ESC differentiation experiment, cell barcodes with more than 1,000 transcripts and fewer than 6,000 genes were selected. Genes detected in fewer than five cells were excluded. Expression levels for each cell were normalized to 10,000 transcripts and logarithmically transformed. Highly variable genes were defined as those with a mean expression value between 0.0125 and 10, and with 0.5 dispersion. The data was then scaled, and the top 4,000 highly variable genes used to generate the UMAP plots. Cells were clustered using the Leiden algorithm.

### Single cell classification and correlation assessment

The three gastrulation datasets (Grosswendt *et al*., Pijuan-Sala *et al*., and Nowotschin *et al*.) were concatenated. Two hundred cells per cell type were randomly selected as the training data. We subset the training data using intersected genes between the training data and the ten CFE studies. Depending on the query data, additional cell types might be added to the training data. The same training data was used for the construction of the SCN, SVM, and SCMAP-cluster meta-gastrulation classifier. SCN meta-gastrulation classifier was constructed with the following parameters: nTopGenes = 30, nTopPairs = 50 after the optimization of parameters shown in Supplementary Figure 3. The default parameters were used for SCMAP and SVM. The assessment metrics of Cohen’s kappa and AUPRC were calculated using assess_comm() function provided by SCN.

### Single-cell gastrulation gene regulatory network enrichment analysis

The meta-gastrulation gene regulatory network was constructed with Context Likelihood of Relatedness (CLR) algorithm from the Grosswendt *et al* gastrulation dataset. Each cell type-specific subnetwork was extracted based on the differentially expressed genes. CLR identified a list of transcription factors and their targets. To compare the *in vivo* and *in vitro* angioblasts and gut endoderm, we subset cells from the *in vivo* and *in vitro* data that were classified by SCN as angioblast and gut endoderm. The subset data were then subjected to the Monocle3 algorithm to correct for batch effect and compute for differentially expressed genes (Trapnell et al., 2014). The differentially expressed genes were used to compute transcription factors, GO terms, and pathways enrichment using the GSEA package (Subramanian et al., 2005).

### Benchmarking training time between SCN (R) and SCN (python)

We used three training datasets, namely Tabula Muris (24,936 cells 52 cell types), Microwellseq (181,755 cells, 125 cell types) and Gastrulation (107,147 cells 48 cell types) to train classifiers separately from both SCN (R) AND SCN (python). We used the default parameters (nTopGenes = 10, nTopPairs = 25) for both pipelines. We bootstrapped the training process 10 times and recorded in the training time for each run. The code for SCN (python) can be found on Github (https://github.com/pcahan1/PySingleCellNet)

### SingleCellNet web application framework construction

To provide easy usage of SCN to a broader range of users, we developed SCN web-application, a user-friendly, interactive online platform that provides classification of user uploaded query samples. SCN web-application was developed using R package shiny (Chang et al., 2021). Interactive visualizations were implemented using R packages ggplot (Wickham, 2016), ggpubr (Kassambara, 2020), and plotly (Sievert, 2020). Front-end UI was developed using R package shinydashboard (Chang and Borges Ribeiro, 2018). Back-end functions for classifying query samples and training classifiers were taken from R package singleCellNet (Tan and Cahan, 2019). To reduce training time, we developed a quick training scheme that skips classification gene selection by generating gene-pairs from the intersecting genes between query samples and pre-selected classification genes in trained classifiers.

## Acknowledgment

This work was supported by the National Institutes of Health under grant R35GM124725 to PC. PC conceived of and supervised the project, performed analysis, and edited the manuscript. YT elaborated on the project idea, performed most of the computational analyses, performed the mouse gastrula experiments, and wrote the manuscript. AS performed all of the embryoid body experiments. MF created pySingleCellNet based on the original R code. DP designed and built the singleCellNet web application. We would like to thank Suraj Kannan, Emily Su, Emily Lo, Bian Qin and Ray Cheng, for providing feedback and support, and additionally to Emily Lo for designing the schematic figure of mouse embryonic development. We also would like to thank Matthew Miyamoto for instruction on the mouse embryo work.

## Supplementary

### Note 1 cluster annotation

| Germ layer | Cluster annotation |
| --- | --- |
| Epiblast | 8: Epiblast |
| Extraembryonic ectoderm and ectoderm lineage | 0: Neural ectoderm anterior<br>1: Neural tube<br>11,0: Extra-embryonic ectoderm<br>11,1: Extra-embryonic ectoderm2<br>20: Surface ectoderm |
| Extraembryonic endoderm and endoderm lineage | 2: Extra-embryonic endoderm<br>12,0: Definitive endoderm<br>18: Extra-embryonic endoderm2<br>19: Parietal endoderm |
| Extraembryonic mesoderm and mesoderm lineage | 3: Neural mesoderm progenitor<br>4: Pre-somitic mesoderm<br>5: Paraxial mesoderm<br>6: Pre-somitic mesoderm2<br>7: Allantois<br>9: Paraxial mesoderm2<br>10: Primitive blood progenitors<br>12,1: Notochord<br>13: Neural mesoderm progenitor2<br>14: Endothelium<br>16: Cardiac progenitor |
| Misc | 15: Unknown1<br>17: Unknown2 |

Cluster 0 is annotated as the neural ectoderm anterior (NEA). Given it emerges at around e7.5 cells, and the expression of Sox2 at mid-late streak stage is restricted at anterior neuroectoderm, which suggests cluster 0 is likely anterior neural ectoderm (Avilion et al., 2003).

Cluster 1 is annotated as the neural tube (NT). The cluster emerges at around E8.25. At late gastrulation, Sox2 expression has extended to the neural tube (Avilion et al., 2003). The high expression of Ptn also suggests neural fate commitment (Li et al., 2017). The Map1b is required for the neural tube formation (Jayachandran et al., 2016). Collectively, they suggest Cluster 1 is the neural tube.

Cluster 2 is annotated as extra-embryonic endoderm (ExEn). Cluster 2 expresses both Ttr and Dab2. Ttr is an endoderm-specific gene that has been expressed in the yolk sac endoderm (Kwon et al., 2008). Dab2 is a pan extra-embryonic endoderm marker (Bérenger). Though cluster 2 has the expression of both Spink1 and Amn, known visceral endoderm markers, we believe that cluster 2 is extra-embryonic endoderm that derives from visceral endoderm.

Cluster 3 is annotated as a neural mesoderm progenitor (NMP) for its expression of Nkx1-2, Cdx2 Cdx1(Rodrigo Albors et al., 2018).

Cluster 4 is annotated as pre-somitic mesoderm (PSM) for its differential expression of Aldh1a2 and Rspo3. Aldh1a indicates cluster 4 might contain somatic cells, while the expression of Rspo3, an important marker for posterior pre-somitic mesoderm (Chal et al., 2018), would suggest this cluster might be mostly pre-somitic.

Cluster 5 is annotated as paraxial mesoderm (PM). Ifitm1 is expressed in the posterior paraxial mesoderm (Tanaka et al., 2005).

Cluster 6 is annotated as pre-somatic mesoderm (PsM) as it expresses Lef1 and Crabp2. Both Lef1 and Crabp1 are involved in Wnt signaling, which has long been indicated to regulate somitogenesis. Lef1 expression has been found in both pre-somitic mesoderm and newly formed somites (Galceran et al., 2004).

Cluster 7 is annotated as allantois (Al) for the expression of Foxf1.

Cluster 8 is annotated as epiblast (Epi), which is composed of primarily e6.5 cells. Cluster 8 has high differentially expressed Fgf5 and Utf1. The Dnmt3b expression in cluster 8 further suggests that cluster 8 is epiblast as its localization in early embryo resembled that of Oct3/4 (Watanabe et al., 2002).

Cluster 9 is annotated as paraxial mesoderm (PM). Prrx2 expressed at pharyngeal arch mesenchyme (de Jong and Meijlink, 1993). Fbn2 expression was detected in paraxial mesoderm in zebrafish (Gansner et al., 2008).

Cluster 10 is annotated as primitive blood progenitors (PBP) for its expression of Tal1, Hbb-bh1, Gata1 (Ibarra-Soria et al., 2018).

Cluster 11,0 and Cluster 11,1 are both annotated as extra-embryonic ectoderm. The top differentially expressed gene in cluster 11,1 is Plet1 which was shown to be highly expressed in mouse placenta, specifically the distal-most part of the extraembryonic ectoderm (Frankenberg et al., 2007; Zhao et al., 2004). Amongst the differentially expressed gene in 11,0, Utf1 was reported to be expressed in extra-embryonic ectoderm and Utf1-mice showed placental insufficiency (Nishimoto et al., 2013). Gjb3 and Rhox5 were also involved in placenta development (Wen et al., 2017).

Cluster 12,0 is annotated as definitive endoderm (DE) which expresses Krt18, Krt8, and Trh. Krt18 was reported to be highly expressed in definitive endoderm (Wen et al., 2017), while Krt8 was reported as highly expressed in visceral endoderm. Additionally, the expression of Trh supports that cluster 12 is more likely to be definitive endoderm than visceral endoderm (McKnight and Hou, 2007).

Cluster 12,1 is annotated as notochord (Nt) for its expression of T and Cthrc1. T is normally associated with notochord development (Stemple, 2005). Cthrc1 is important for regulating notochord elongation and cell migration during gastrulation (Cheng et al., 2019).

Cluster 13 is annotated as neural mesoderm progenitor (NMP) for its expression of Nkx1-2, Cdx2, Cdx1.

Cluster 14 is annotated as endothelium (Endo) for its expression of Fli-1, LMO2, and Kdr. Fli-1 was shown to bind to the promoter of endothelial cells in vivo (Pimanda et al., 2006). Lmo2 is required for yolk sac erythropoiesis. Kdr is known as vascular endothelial growth factor receptor 2 (Zoldan and Lytton).

Cluster 15 is annotated as unknown. Prdx1 is important for vascular development (Prdx1-encoded peroxiredoxin is important for vascular development in zebrafish). However, the expression of Gapdh, Tuba1b, Tubb5, and Eif4a1 in cluster 15 might suggest that the cells in this cluster might be damaged. We decided to remove this cluster from further analysis.

Cluster 16 is annotated as an early cardiac progenitor (CP). Csrp1 is expressed in the endoderm underlying cardiac mesoderm and a restricted region of cardiac mesoderm (Miyasaka et al., 2007). Tpm1 is strongly expressed in the presomitic mesoderm (Chal et al., 2018). On E8.5, Dok4 was found to be preferentially expressed in developing endocardial cells (Narumiya et al., 2007).Cdkn1c is an imprinted gene that primarily expresses in the placenta (Singh et al., 2017). In addition to neural crest cells, Cnn2 also expresses in the developing heart (Ulmer et al., 2013). Smarcd3 is a marker of early cardiac progenitor which expresses before the known cardiac progenitor marker (Nkx2-5, Isl1, and Tbx5), and it lies at the anterior-proximal region of the embryo and extends into the extraembryonic tissue (Devine et al., 2014).

Cluster 17 is also annotated as unknown. The gene expression profile of cluster 17 is similar to that of cluster 15, which might also suggest damaged or stressed cells. We decided to remove this cluster from further analysis.

Cluster 18 is annotated as extra-embryonic endoderm (ExEn) for its expression of Ttr and Dab2.

Cluster 19 is annotated as parietal endoderm (ParE) for its expression of Srgn and Lama1. Lama1 is a known marker for parietal endoderm (Miner et al., 2004).

Cluster 20 is annotated as surface ectoderm (SE) for its expression of Wnt6 and Slc2a3. Wnt6 expresses in surface ectoderm during gastrulation (Geetha-Loganathan et al., 2006). Slc2a3 expression has been detected in surface ectoderm at late gastrula (E8.5) (Sousa-Nunes et al., 2003).

#### Box 1 Abbreviation used for cell types

|  | abbreviation | Cell type |
| --- | --- | --- |
| 1 | AI | Allantois |
| 2 | AmE | Amniotic ectoderm |
| 3 | AngB | Angioblast |
| 4 | APS | Anterior primitive streak |
| 5 | Br | Forebrain & rostral hindbrain |
| 6 | DE | Definitive endoderm |
| 7 | Ect | Ectoderm |
| 8 | emVE | Embryonic visceral endoderm |
| 9 | EpC | Ectoplacental cone |
| 10 | EPI | Epiblast |
| 11 | ExE | Extra-embryonic ectoderm |
| 12 | exVE | Extra-embryonic visceral endoderm |
| 13 | FHF | First heart field |
| 14 | GE | Gut endoderm |
| 15 | HemaB | Hemangioblast |
| 16 | HSC | Hematopoietic stem cell |
| 17 | ICM | Inner cell mass |
| 18 | InM | Intermediate mesoderm |
| 19 | LPM | Lateral plate mesoderm |
| 20 | mTE | mural trophectoderm |
| 21 | NC | Neural crest |
| 22 | NP | Neural plate |
| 23 | NPB | Neural plate border |
| 24 | NT | Neural tube |
| 25 | Nt | Notochord |
| 26 | ParE | Parietal endoderm |
| 27 | PhM | Pharyngeal mesoderm |
| 28 | PHT | Primitive heart tube |
| 29 | PM | Paraxial mesoderm |
| 30 | PpE | Preplacodal ectoderm |
| 31 | PPS | Posterior primitive streak |
| 32 | PrE | Primitive endoderm |
| 33 | PrN | Primitive node |
| 34 | PsM | Presomatic mesoderm |
| 35 | pTE | polar trophectoderm |
| 36 | SHF | Second heart field |
| 37 | Smt | Somites |
| 38 | SoM | Somitic mesoderm |
| 39 | SpM | Splanchnic mesoderm |
| 40 | SuE | Surface ectoderm |
| 41 | TE | Trophectoderm |
| 42 | TSC | Trophoblast stem cell |
| 43 | VE | Visceral endoderm |

### Note 2 Details on biomimetic studies

We only obtained one time point (120h or D5) for the gastruloid study and gathered a wide range of time points for the EB studies. There is only one scRNA-seq study on gastruloid (van den Brink et al., 2020). Mouse ESCs were cultured in N2B27 medium. When mESCs aggregates were detected at around d2 to d4, a pulse of Wnt agonist CHIR99021 (Chi) was given.

EB_multiseq was generated in this study as described in Figure 3. Brachyury-GFP ESCs were induced with two primitive streak (PS) induction treatments (WAB and WAG) on day 2 for 48 hours, followed by the bFGF treatment on day 4.

EB_replicate and EB_recording are two scRNA-seq experiments from Kim *et al* study (Kim et al., 2020). EB_replicate is a collection of 1,281 cells from all stages of the EB time course (D0-D14). In the absence of LIF, the mESCs were spontaneous differentiated and aggregated into EB over a course of 14 days. Single cells were sampled at a 48hr interval. EB_recording is a collection of 3,489 cells with 14 days of in vitro culture. In this experiment, ten tandem LoxP sites with a static barcode of ten random nucleotides, which were termed the unique clonal identifiers (UCIs), were introduced to mESCs to trace unique clonal lineage. EB_recording contains day 14 cells where tandem-loxP barcodes were induced at two time points: ESCs (day 0) and after expression of postimplantation epiblast marker genes (days 8/9). The barcode information was used in the original paper for lineage tracing; However, we did not filter cells based on that criterion.

**Supplementary Figure 1 Depiction of early mouse development through gastrulation**

**Supplementary Figure 2 Batch correction forces similarity between engineered cells and reference cell types** (**A**) Three gastrulation scRNA-seq *in vivo* references datasets and one query directed differentiation study (Spangler et al., 2018) were integrated together with Canonical Correlated Analysis in Seurat. (left) The UMAP embeddings of all cells colored by batch (right) The UMAP embeddings of all cells colored by cell type annotations provided by the authors (**B**) (left) UMAP embeddings of all *in vivo* gastrulation reference cell types and day 4 of the query dataset (right) UMAP embeddings of all *in vivo* gastrulation reference cell types and day 6 of the query dataset. (C) The same UMAP embeddings as B, but cells are colored by classification score of notochord and epiblast

**Supplementary Figure 3 Parameter optimizations for SCN meta-gastrulation classifier** (**A**) A scatter plot of performance of SCN classifiers with different combinations of parameters. (**B**) The PR curves of the optimized classifier for each cell type (**C**) The classification heatmap of the heldout data for the meta-gastrulation classifier.

**Supplementary Figure 4 Quality control and raw data for the mouse gastrulation scRNA-seq data** (**A**) Number of cells sequenced per gastrulation timepoint (**B**-**C**) Median UMI counts and median gene counts per cell for each time point (**D**) Experimental details of how the scRNA-seq for each gastrulation time point was generated

**Supplementary Figure 5 SCN classification of Spangler et al 2018 study** (**A**) (left) A UMAP embeddings for the day 6 direct differentiated cells colored by clusters (right) A UMAP embeddings for the day 4 direct differentiated cells colored by clusters (**B**) The classification heatmap for both day 4 and day 6 with the meta-gastrulation classifier. x-axis is the SCN classifier categories, y axis contains a single cell grouped by clusters. The SCN classification score ranges from 0 to 1.

**Supplementary Figure 6 Quality control and additional figures for the EB scRNA-seq study** (**A**) median UMI counts for each PS induction methods that are sorted and unsorted (**B**) Total number of cells for each treatment and sorting conditions (**C**) A UMAP embedding of EB scRNA-seq profiles colored by SCN classification score for epiblast, inner cell mass and trophoblast stem cell. (**D**) A complete SCN classification heatmap for all EB scRNA-seq profiles from the meta-gastrulation classifier. (**E**) SCN classification for sorted WAB treated cells (**F**) SCN classification for unsorted WAB treated cells (**G**) SCN classification for sorted WAG treated cells (**H**) SCN classification for unsorted WAG treated cells

**Supplementary Figure 7 Correlation based *in vitro* and *in vivo* timepoints alignment** The cell type composition for each time point was calculated from SCN analysis. Then *in vitro* and *in vivo* cell type similarity for each time point was calculated with cosine similarity. (**A**) The cosine similarity-based time point alignment between EB_multiseq and Grosswendt*et al* gastrulating embryos. (**B**) The cosine similarity-based time point alignment between EB_recording and Grosswendt*et al* gastrulating embryos. (**C**) The cosine similarity-based time point alignment between EB_replicate and Grosswendt*et al* gastrulating embryos. (**D**) The cosine similarity-based time point alignment between Gastruloid and Grosswendt *et al* gastrulating embryos

**Supplementary Figure 8 Extended comparisons for the biomimetic studies** (**A**-**O**) Distribution and violin plots for the classification score for each cell type colored by each individual study

**Supplementary Figure 9 Gene Set enrichment analysis for angioblasts and gut endoderm** Using *in vivo* cell types as references, the differentially expressed genes (DEGs) for each *in vitro* study were calculated from Monocle3. The DEGs were then used in this gene set enrichment analysis (GSEA). (**A**-**B**) GSEA for EB_recording angioblasts on MSigDB C2 and C5 gene set collections. (**C**-**D**) GSEA for EB_multiseq gut endoderm on MSigDB C2 and C5 gene set collections.

**Supplementary Figure 10 Single-cell network enrichment analysis of angioblasts and gut endoderm** Single-cell network enrichment analysis for angioblasts **(A-C)** and gut endoderm **(D-F)** was performed to identify differentially enriched transcription factor networks. (**A**) Top 20 positively and negatively enriched transcription factor networks for EB_multiseq angioblasts. (**B**) Top 20 positively and negatively enriched transcription factor networks for EB_replicate angioblasts. (**C**) Top 20 positively and negatively enriched transcription factor networks for Gastruloid angioblasts. (**D**) Top 20 positively and negatively enriched transcription factor networks for EB_recording gut endoderm. (**D**) Top 20 positively and negatively enriched transcription factor networks for EB_replicate gut endoderm. (**E**) Top 20 positively and negatively enriched transcription factor networks for Gastruloid gut endoderm.

