## Supplementary Materials for "Quantitative comparison of *in vitro and in vivo* embryogenesis at a single cell resolution"

### Supplementary Figure 1

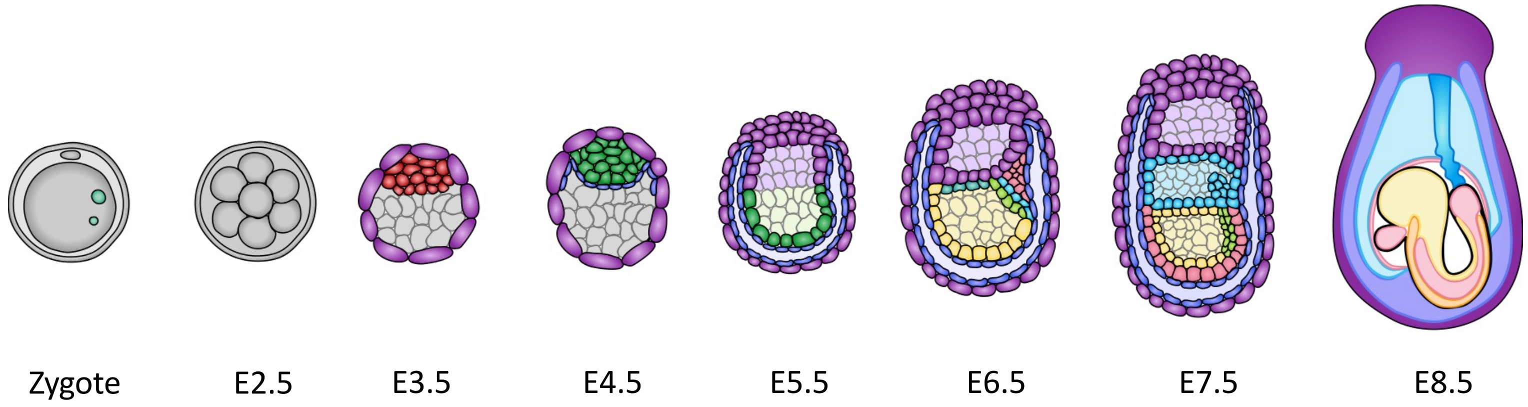

### Supplementary Figure 2

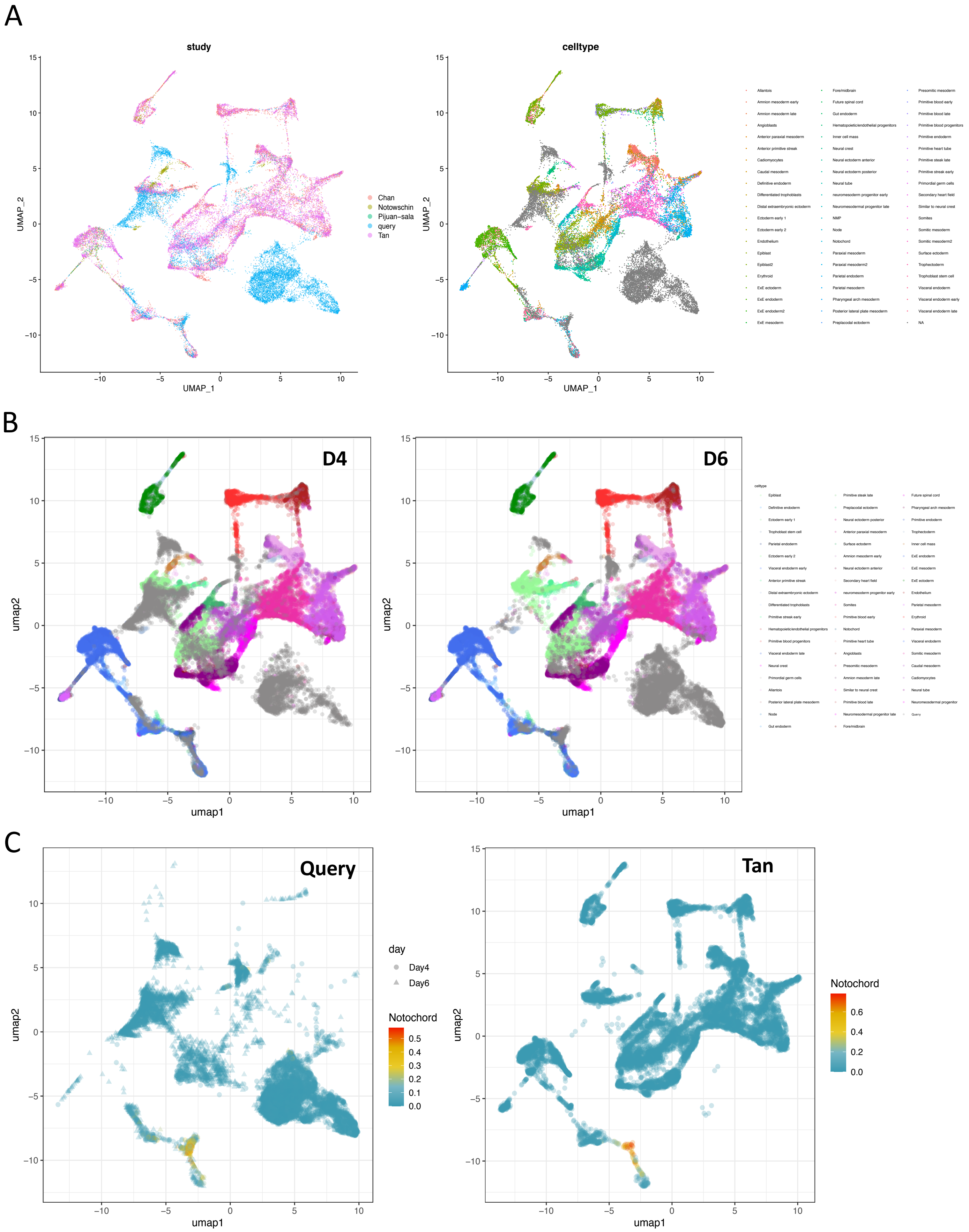

D

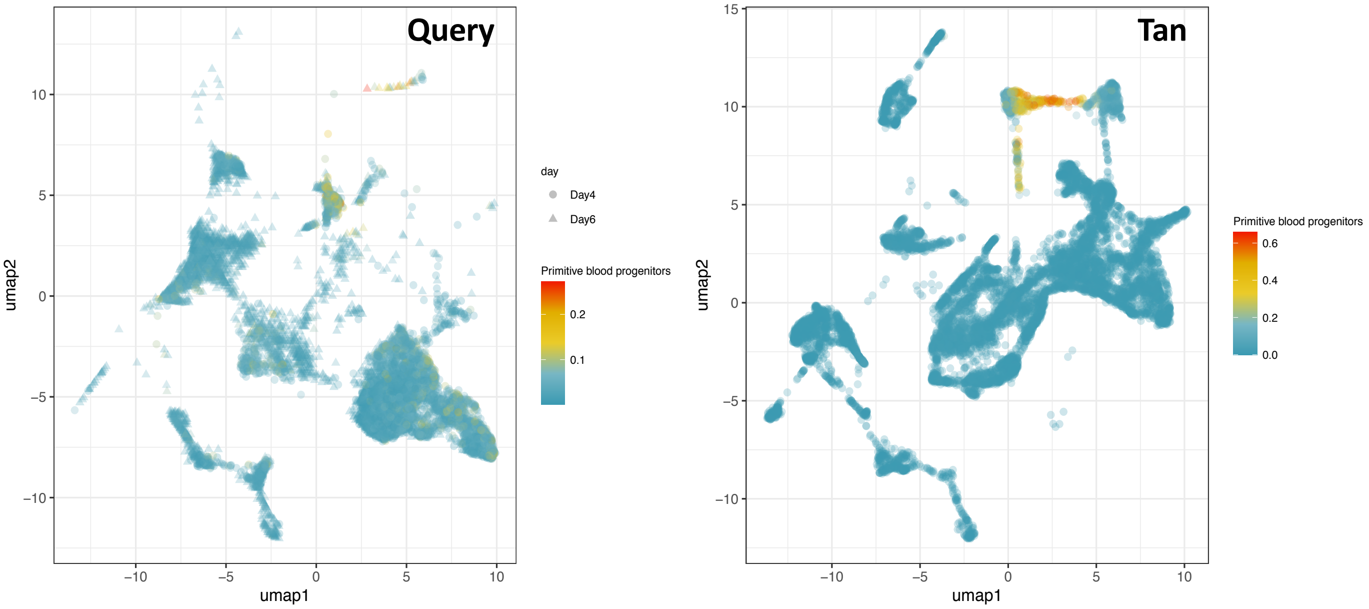
Query
Tan
umap1
umap2
day
Day4
Day6
Primitive blood progenitors
0.2
0.1
0.6
0.4
0.2
0.0

### Supplementary Figure 3

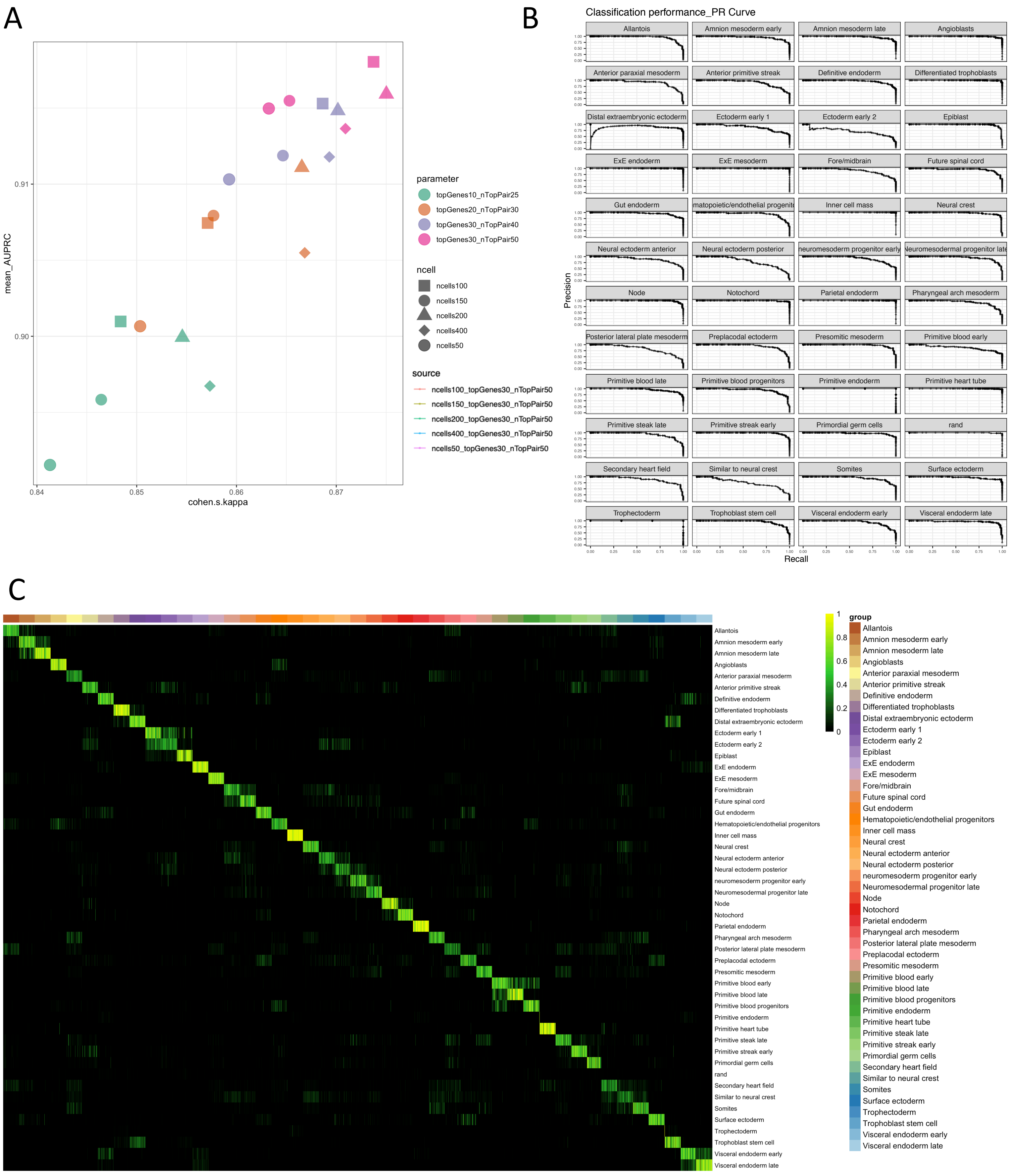

### Supplementary Figure 4

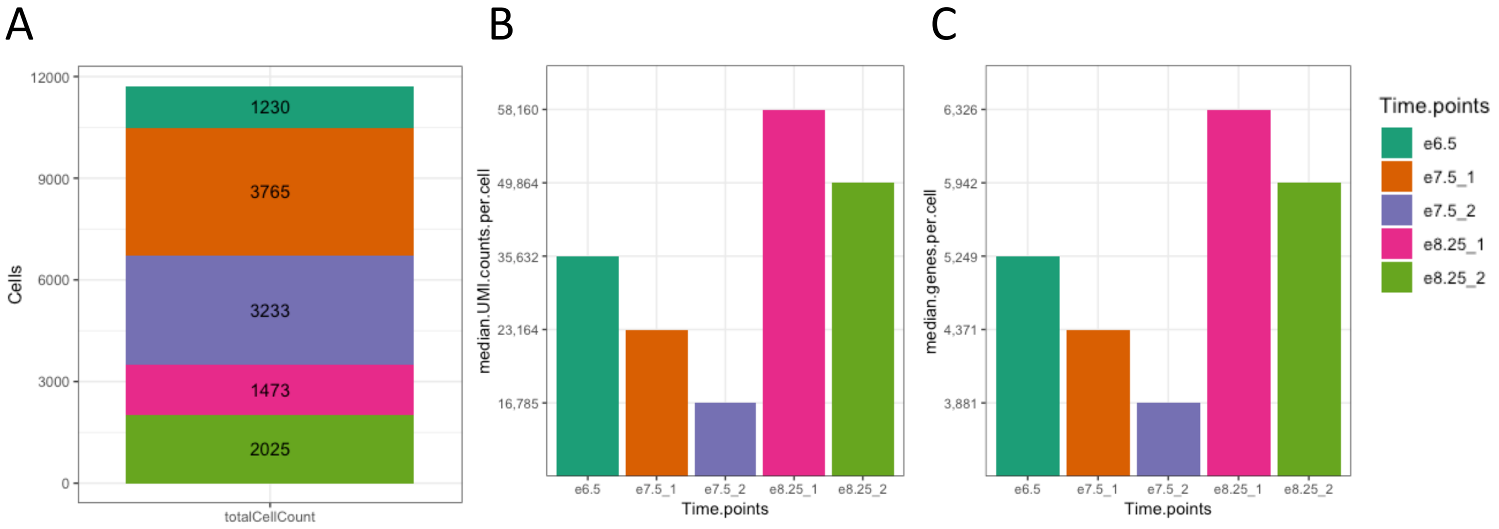


D

| Date of experiment | Time points | Emrbyos | Cells | Seq | mean reads per cell | median genes per cell | median UMI counts per cell | Time_point |
|---|---|---|---|---|---|---|---|---|
| **8/23/18** | e6.5 | 13 | 1230 | NovaSeq | 363,452 | 5,249 | 35,632 | e6.5 |
| **9/6/18** | e7.5_1 | 8 | 3765 | NovaSeq | 86,103 | 4,371 | 23,164 | e7.5 |
| **9/6/18** | e7.5_2 | 8 | 3233 | NovaSeq | 141,595 | 3,881 | 16,785 | e7.5 |
| **4/5/18** | e8.25_1 | 1 | 1473 | NovaSeq | 284,417 | 6,326 | 58,160 | e8.25 |
| **4/5/18** | e8.25_2 | 1 | 2025 | NovaSeq | 218,973 | 5,942 | 49,864 | e8.25 |

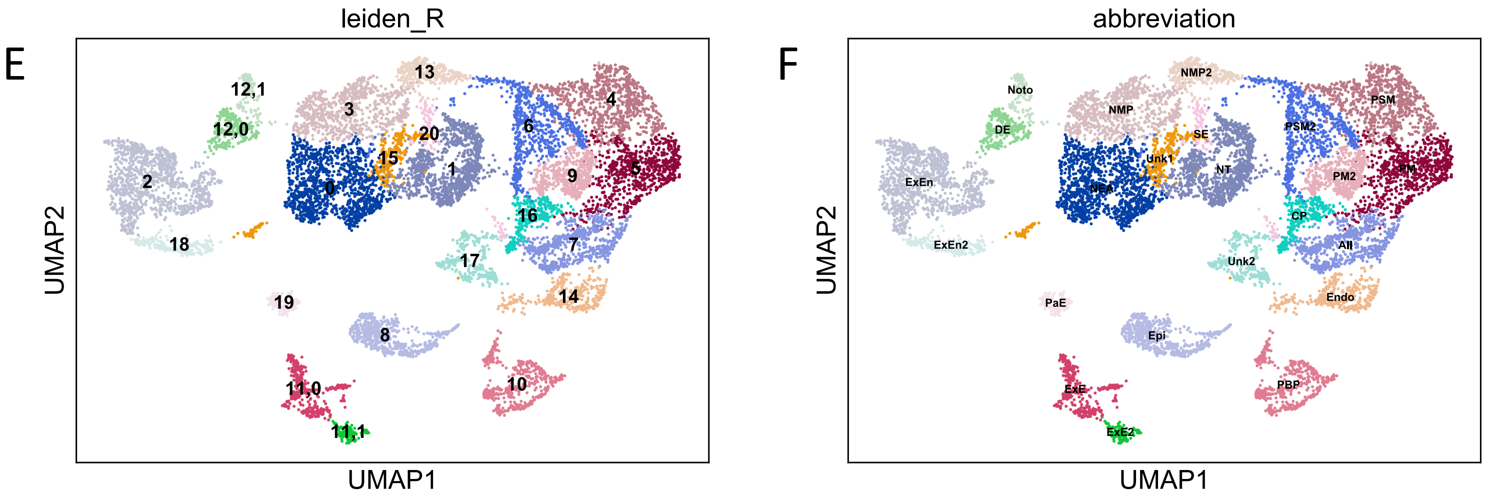

G

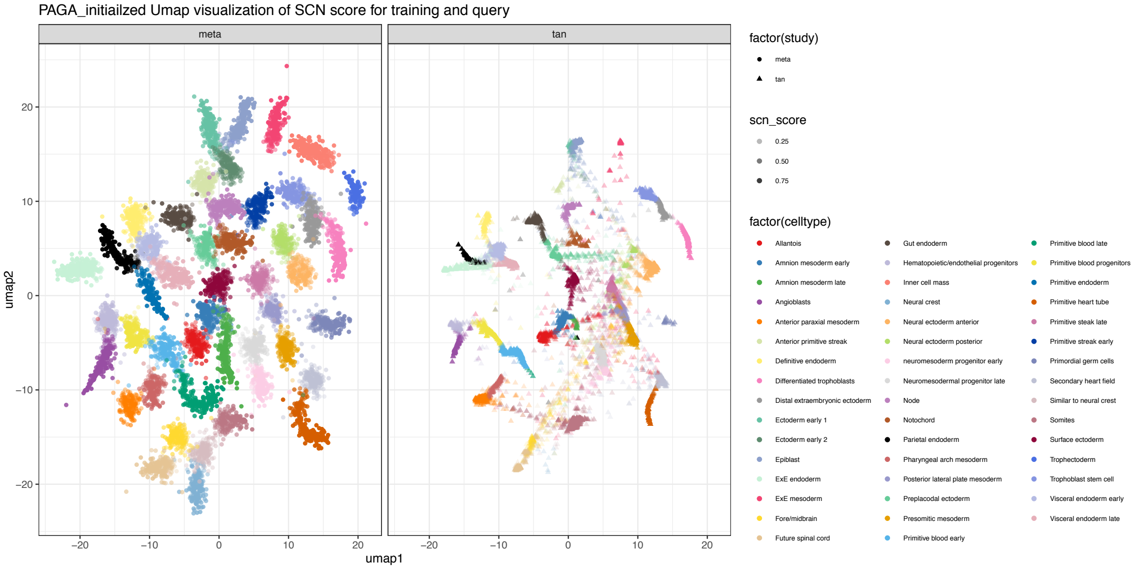
PAGA_initiailzed Umap visualization of SCN score for training and query
meta
tan
umap1
umap2
factor(study)
meta
tan
scn_score
0.25
0.50
0.75
factor(celltype)
Allantois
Amnion mesoderm early
Amnion mesoderm late
Angioblasts
Anterior paraxial mesoderm
Anterior primitive streak
Definitive endoderm
Differentiated trophoblasts
Distal extraembryonic ectoderm
Ectoderm early 1
Ectoderm early 2
Epiblast
ExE endoderm
ExE mesoderm
Fore/midbrain
Future spinal cord
Gut endoderm
Hematopoietic/endothelial progenitors
Inner cell mass
Neural crest
Neural ectoderm anterior
Neural ectoderm posterior
neuromesoderm progenitor early
Neuromesodermal progenitor late
Node
Notochord
Parietal endoderm
Pharyngeal arch mesoderm
Posterior lateral plate mesoderm
Preplacodal ectoderm
Presomitic mesoderm
Primitive blood early
Primitive blood late
Primitive blood progenitors
Primitive endoderm
Primitive heart tube
Primitive steak late
Primitive streak early
Primordial germ cells
Secondary heart field
Similar to neural crest
Somites
Surface ectoderm
Trophectoderm
Trophoblast stem cell
Visceral endoderm early
Visceral endoderm late

H

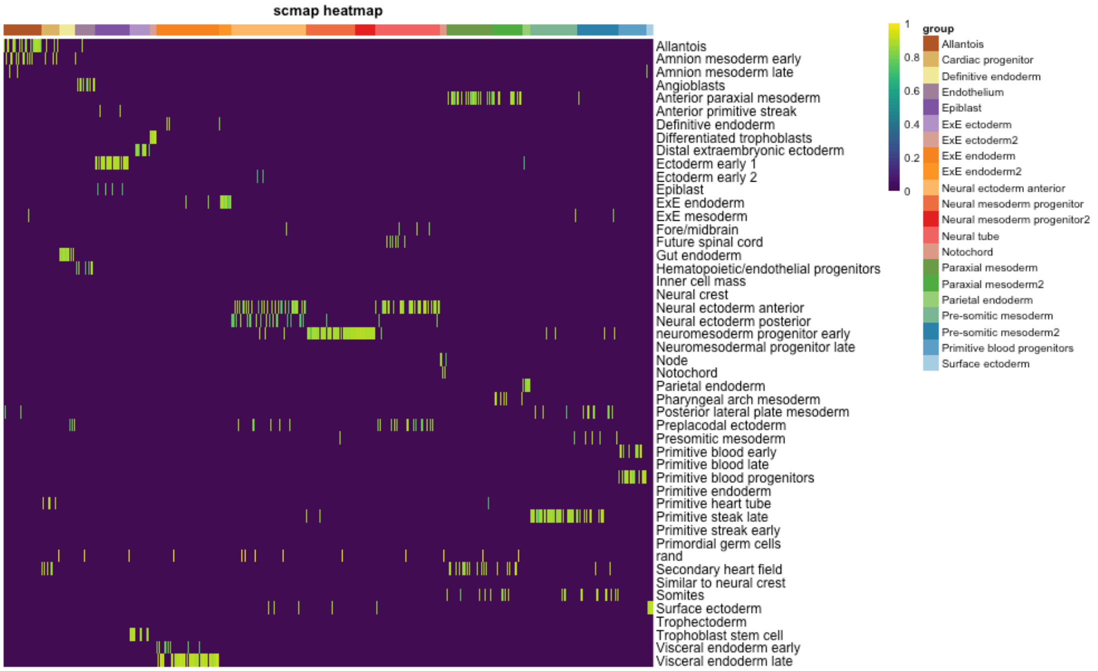
scmap heatmap
Allantois
Amnion mesoderm early
Amnion mesoderm late
Angioblasts
Anterior paraxial mesoderm
Anterior primitive streak
Definitive endoderm
Differentiated trophoblasts
Distal extraembryonic ectoderm
Ectoderm early 1
Ectoderm early 2
Epiblast
ExE endoderm
ExE mesoderm
Fore/midbrain
Future spinal cord
Gut endoderm
Hematopoietic/endothelial progenitors
Inner cell mass
Neural crest
Neural ectoderm anterior
Neural ectoderm posterior
neuromesoderm progenitor early
Neuromesodermal progenitor late
Node
Notochord
Parietal endoderm
Pharyngeal arch mesoderm
Posterior lateral plate mesoderm
Preplacodal ectoderm
Presomitic mesoderm
Primitive blood early
Primitive blood late
Primitive blood progenitors
Primitive endoderm
Primitive heart tube
Primitive steak late
Primitive streak early
Primordial germ cells
rand
Secondary heart field
Similar to neural crest
Somites
Surface ectoderm
Trophectoderm
Trophoblast stem cell
Visceral endoderm early
Visceral endoderm late
group
Allantois
Cardiac progenitor
Definitive endoderm
Endothelium
Epiblast
ExE ectoderm
ExE ectoderm2
ExE endoderm
ExE endoderm2
Neural ectoderm anterior
Neural mesoderm progenitor
Neural mesoderm progenitor2
Neural tube
Notochord
Paraxial mesoderm
Paraxial mesoderm2
Parietal endoderm
Pre-somitic mesoderm
Pre-somitic mesoderm2
Primitive blood progenitors
Surface ectoderm

I

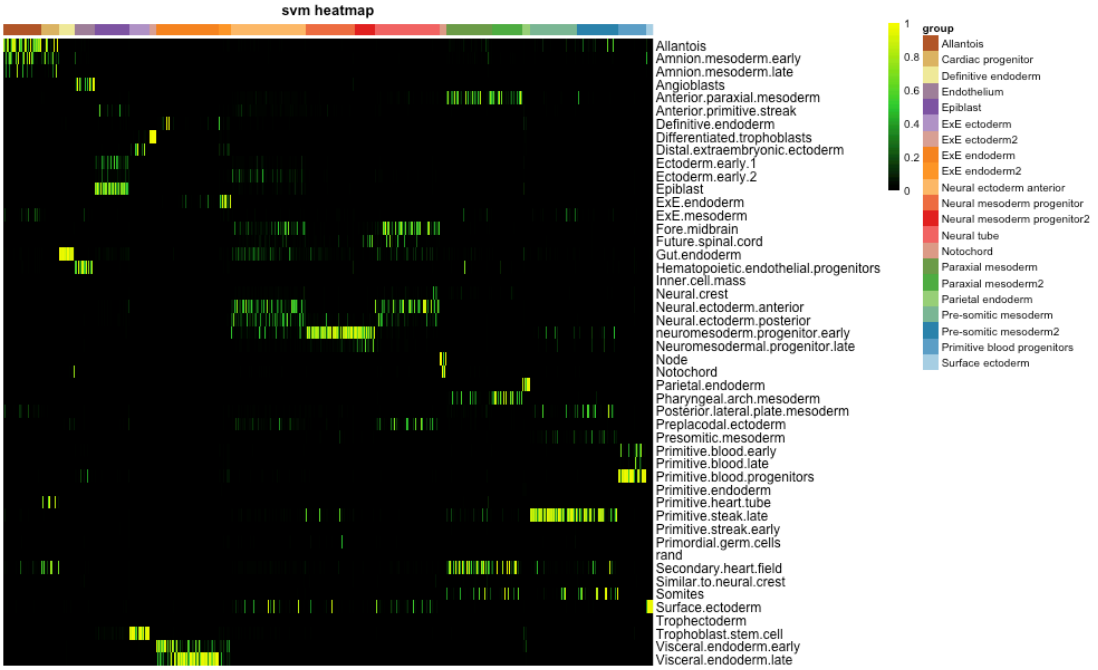
svm heatmap
Allantois
Amnion.mesoderm.early
Amnion.mesoderm.late
Angioblasts
Anterior.paraxial.mesoderm
Anterior.primitive.streak
Definitive.endoderm
Differentiated.trophoblasts
Distal.extraembryonic.ectoderm
Ectoderm.early.1
Ectoderm.early.2
Epiblast
ExE.endoderm
ExE.mesoderm
Fore.midbrain
Future.spinal.cord
Gut.endoderm
Hematopoietic.endothelial.progenitors
Inner.cell.mass
Neural.crest
Neural.ectoderm.anterior
Neural.ectoderm.posterior
neuromesoderm.progenitor.early
Neuromesodermal.progenitor.late
Node
Notochord
Parietal.endoderm
Pharyngeal.arch.mesoderm
Posterior.lateral.plate.mesoderm
Preplacodal.ectoderm
Presomitic.mesoderm
Primitive.blood.early
Primitive.blood.late
Primitive.blood.progenitors
Primitive.endoderm
Primitive.heart.tube
Primitive.steak.late
Primitive.streak.early
Primordial.germ.cells
rand
Secondary.heart.field
Similar.to.neural.crest
Somites
Surface.ectoderm
Trophectoderm
Trophoblast.stem.cell
Visceral.endoderm.early
Visceral.endoderm.late
group
Allantois
Cardiac progenitor
Definitive endoderm
Endothelium
Epiblast
ExE ectoderm
ExE ectoderm2
ExE endoderm
ExE endoderm2
Neural ectoderm anterior
Neural mesoderm progenitor
Neural mesoderm progenitor2
Neural tube
Notochord
Paraxial mesoderm
Paraxial mesoderm2
Parietal endoderm
Pre-somitic mesoderm
Pre-somitic mesoderm2
Primitive blood progenitors
Surface ectoderm

### Supplementary Figure 5

A

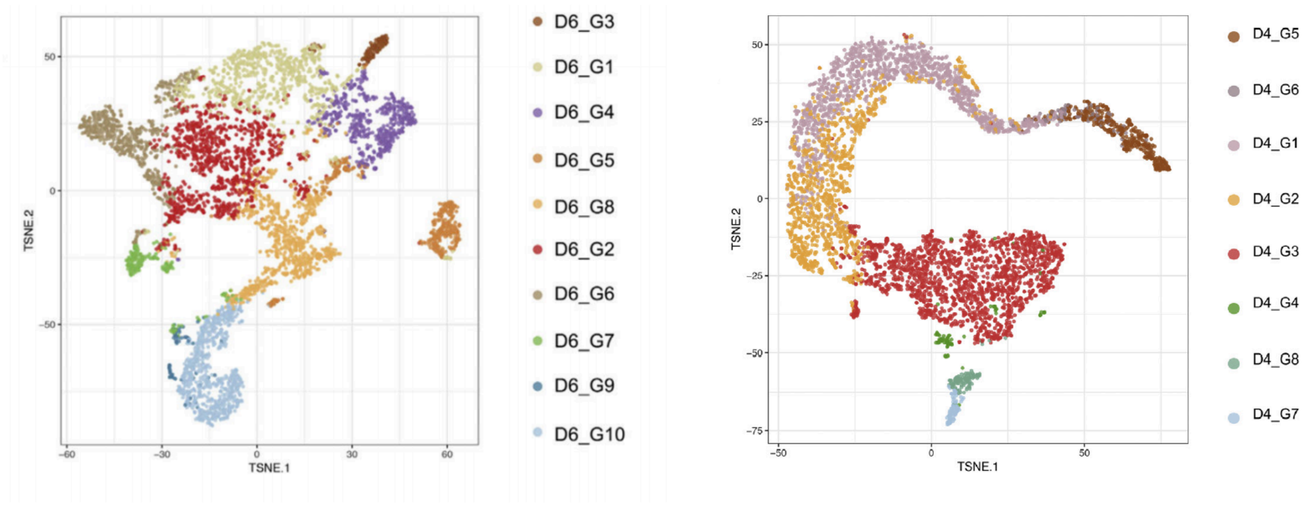


B

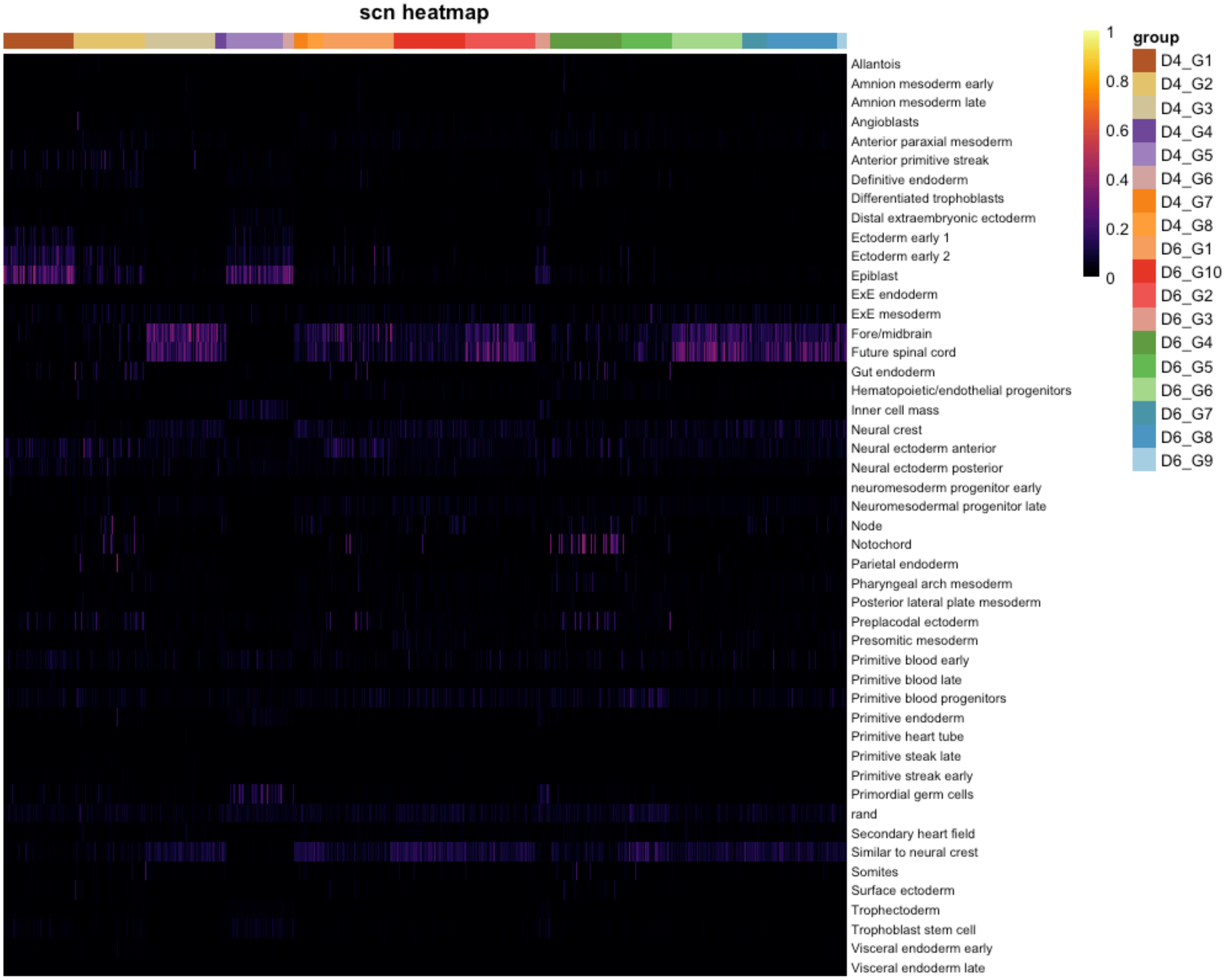

#### Supplementary Figure 6

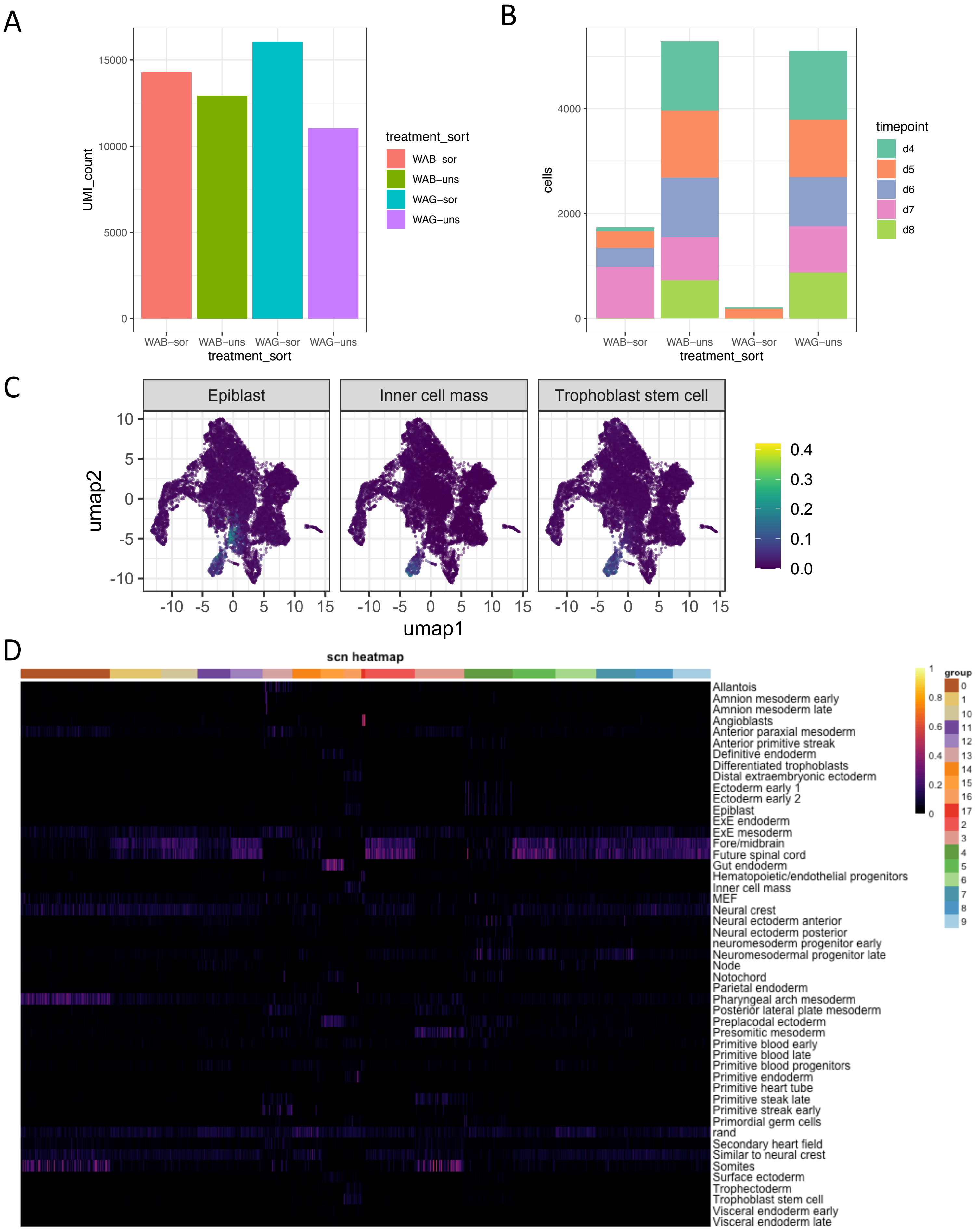

E SCN classification for sorted WAB treated cells

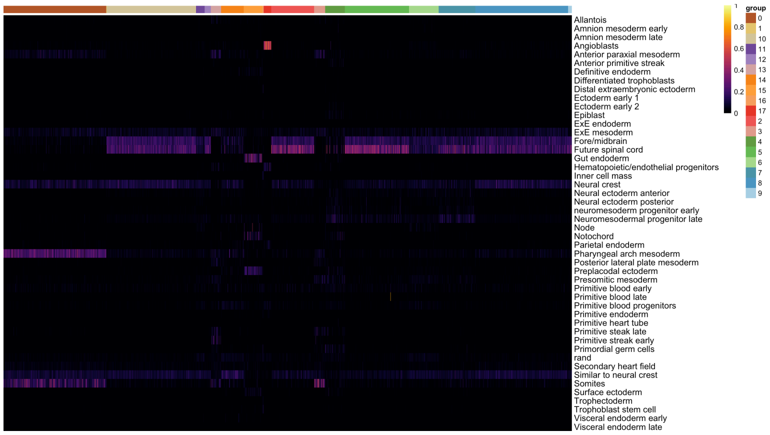


F SCN classification for unsorted WAB treated cells

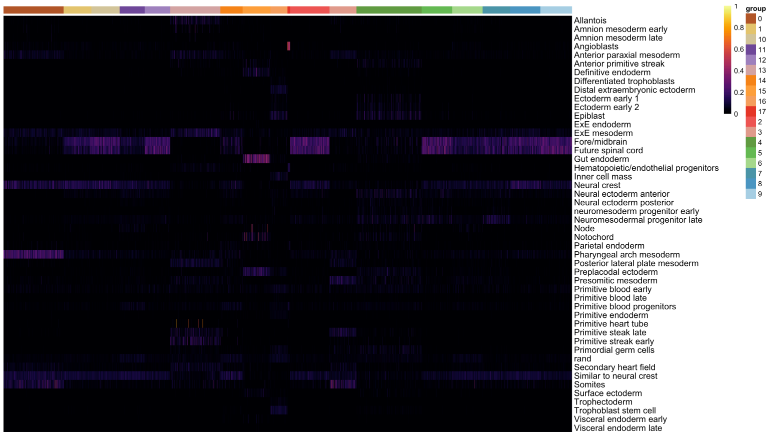


G SCN classification for sorted WAG treated cells

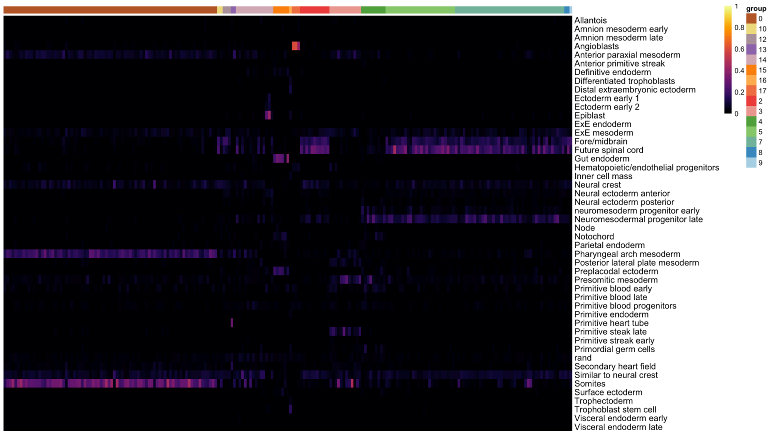


H SCN classification for unsorted WAG treated cells

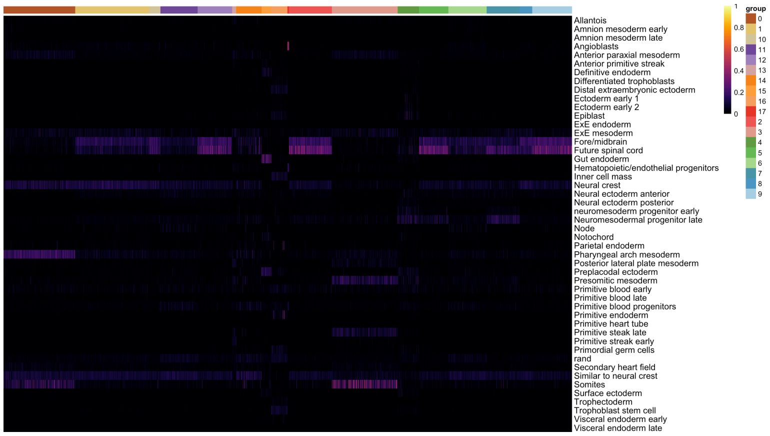

### Supplementary Figure 7

A

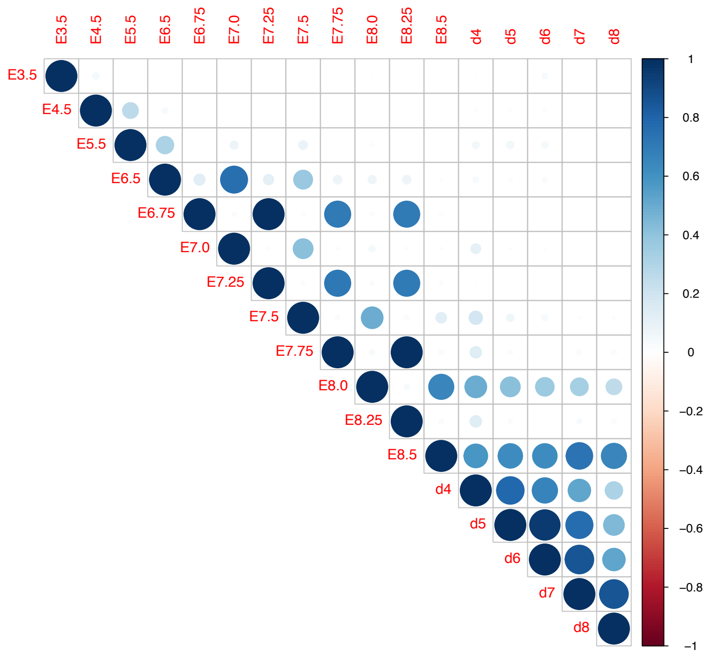


B

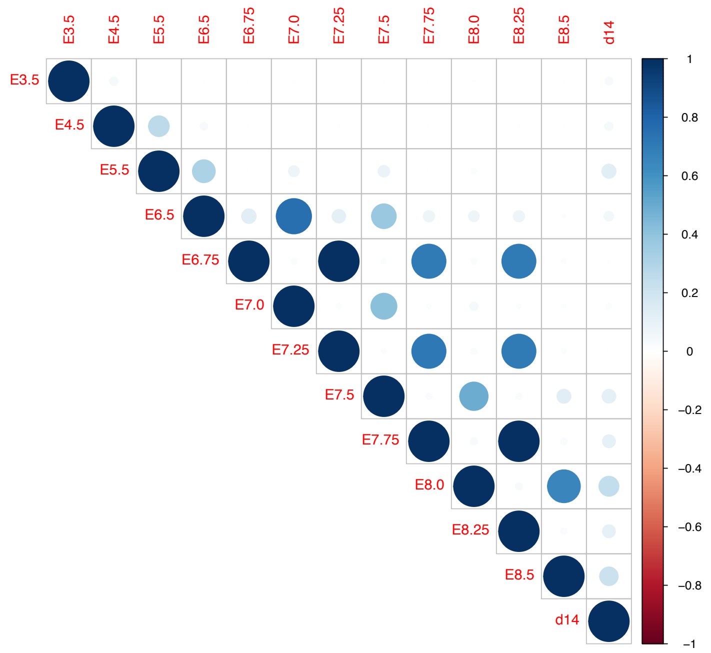


C

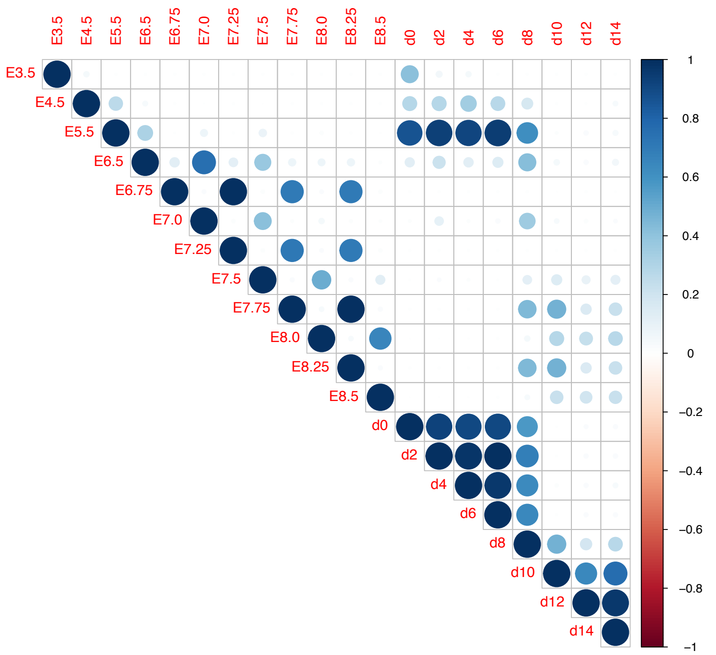


D

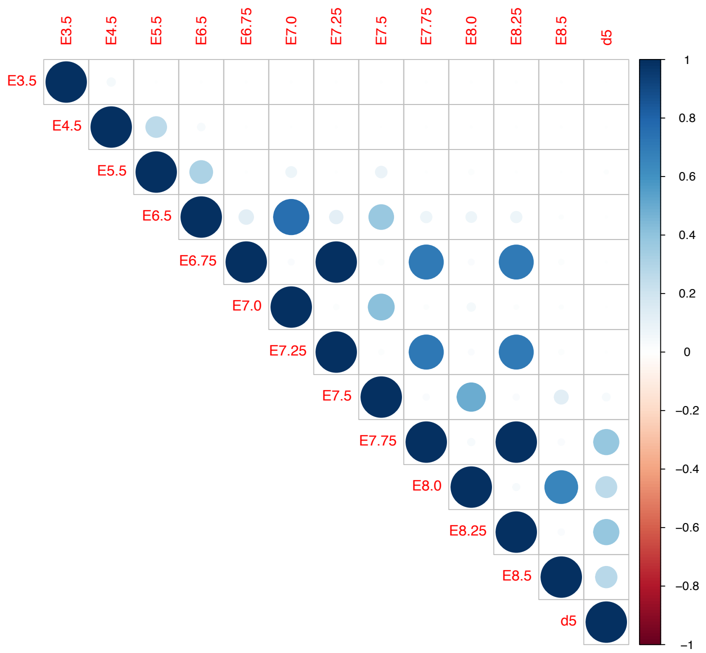

### Supplementary Figure 8

A

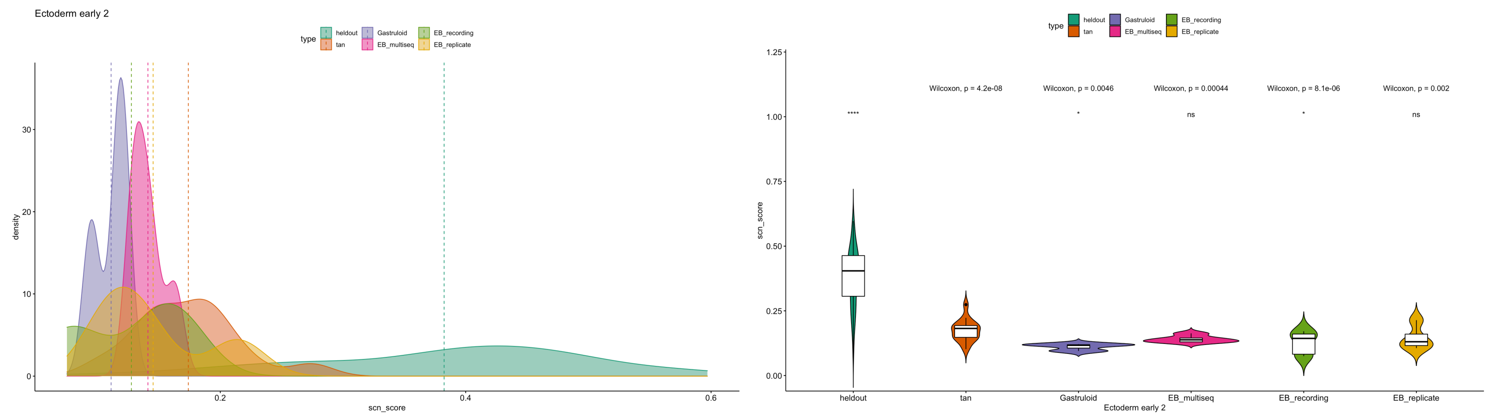


B

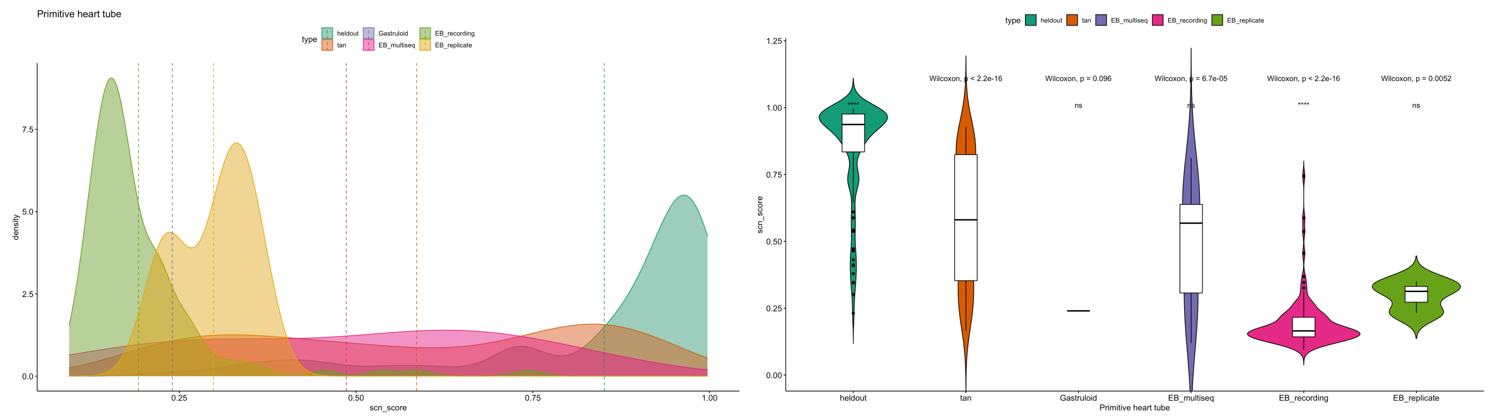


C

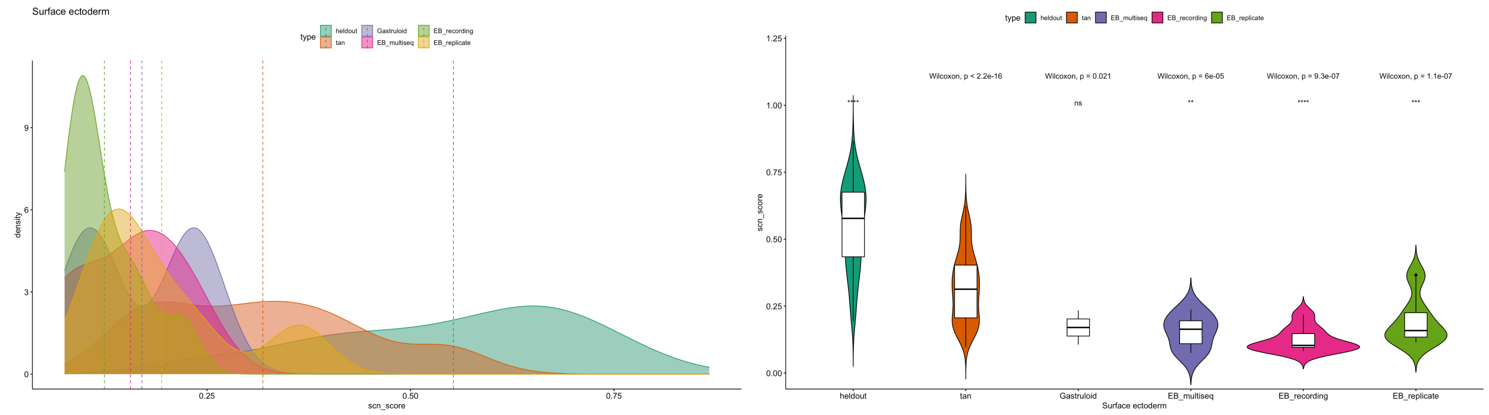


D

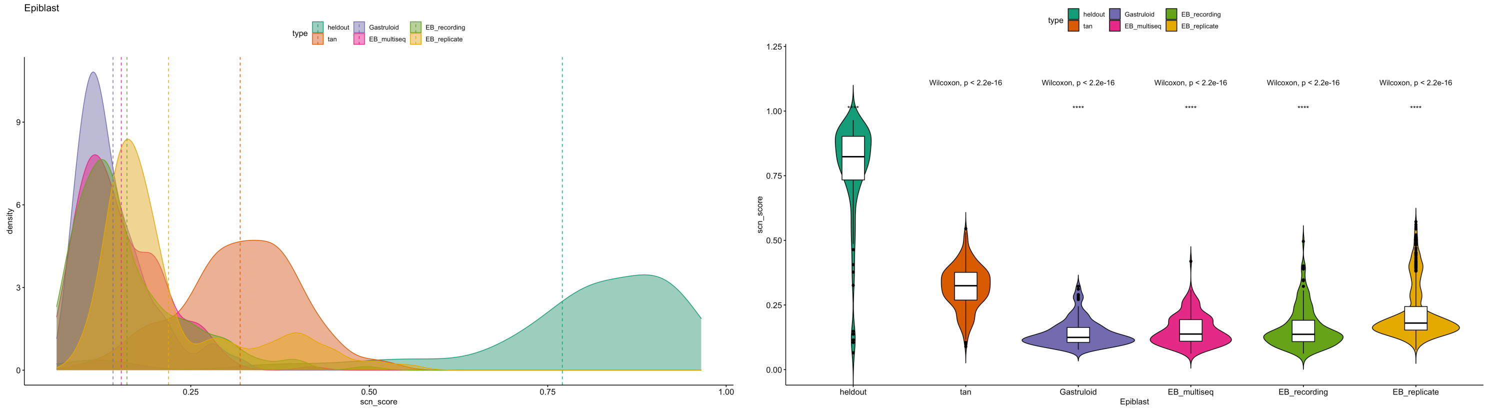

### Supplementary Figure 8

E

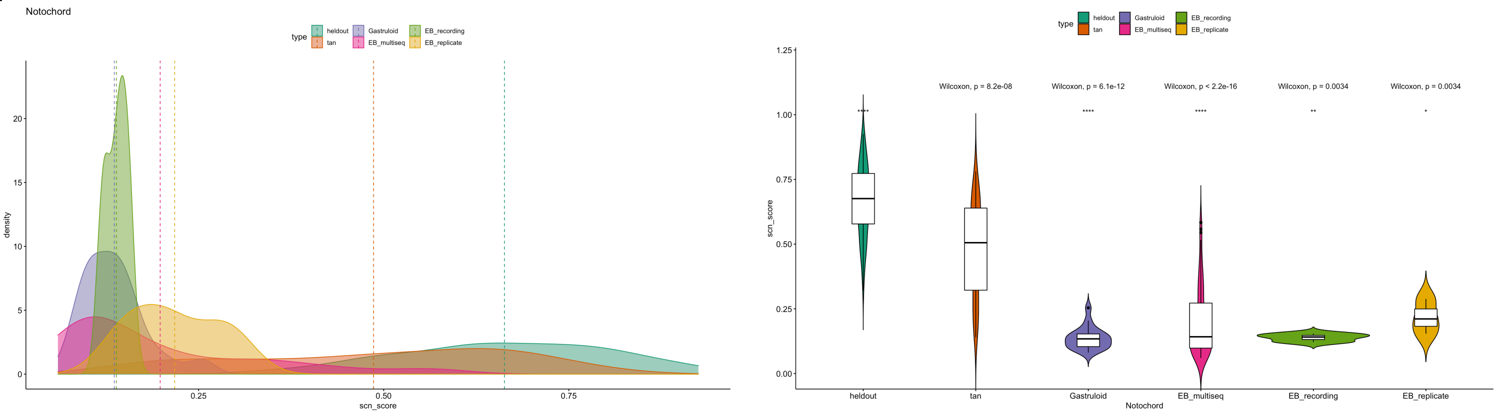


F

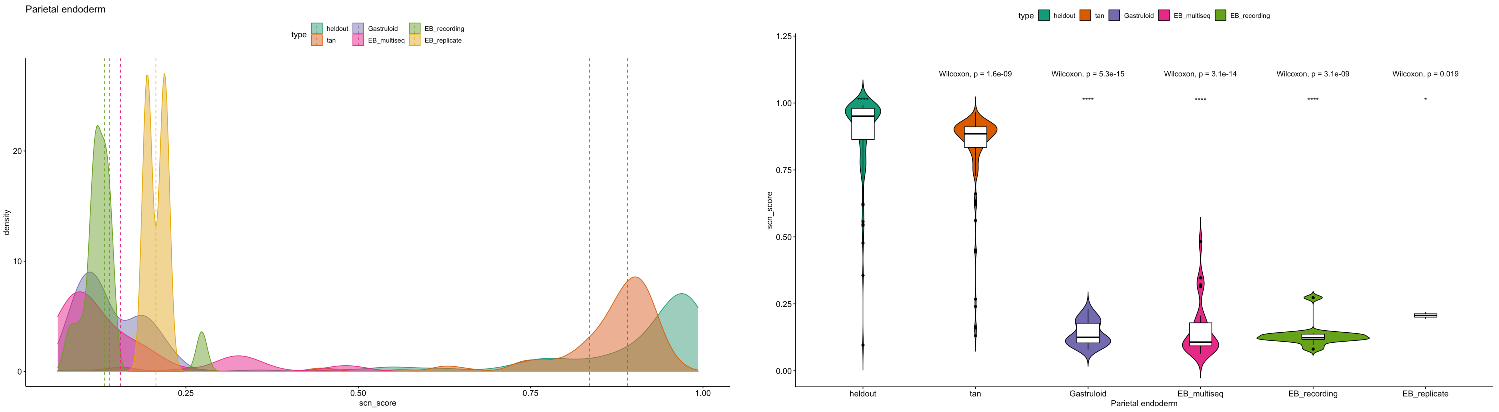


G

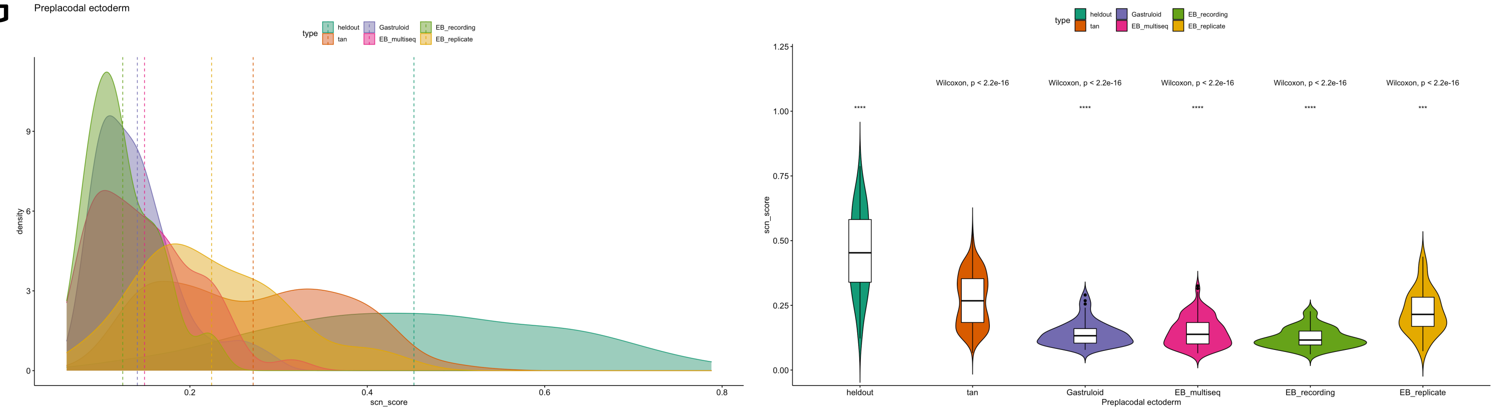


H

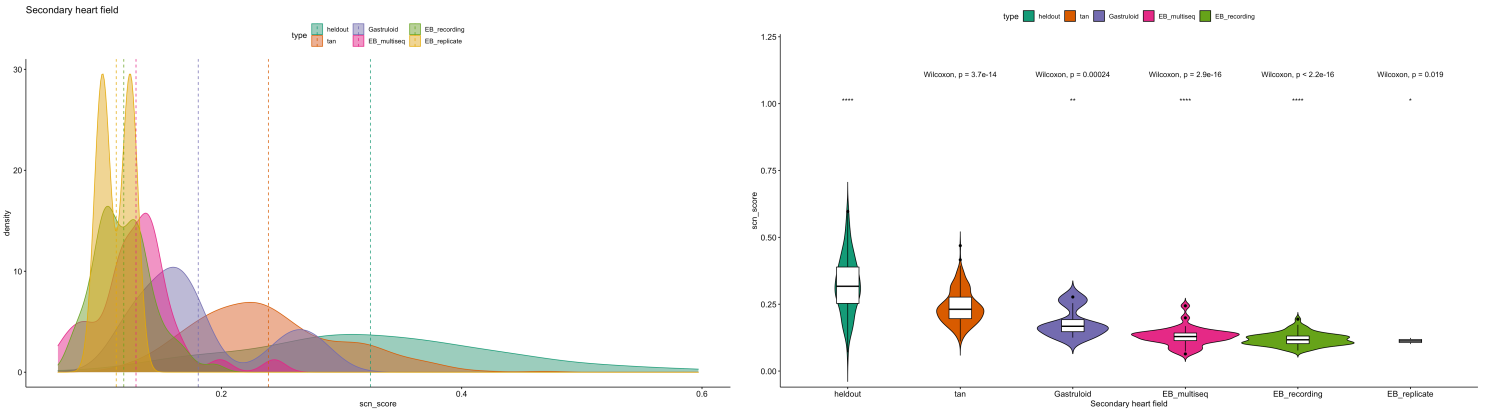

### Supplementary Figure 8

I

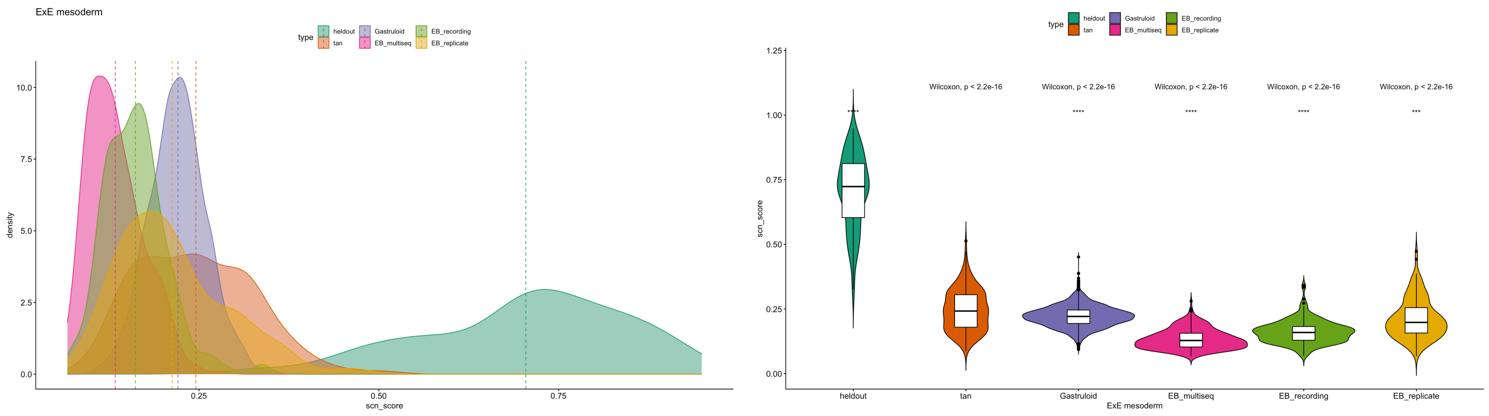


J

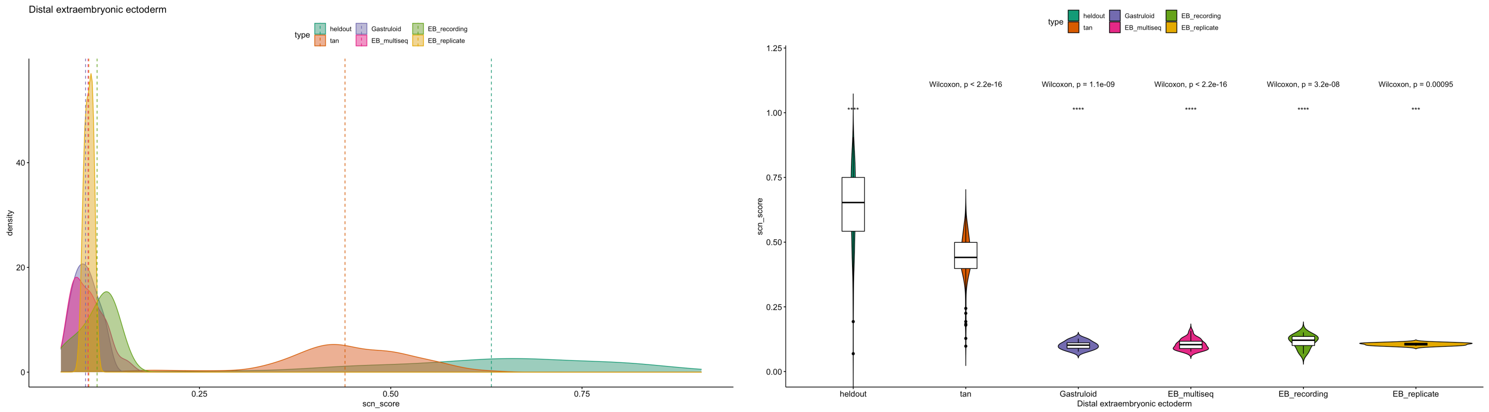


K

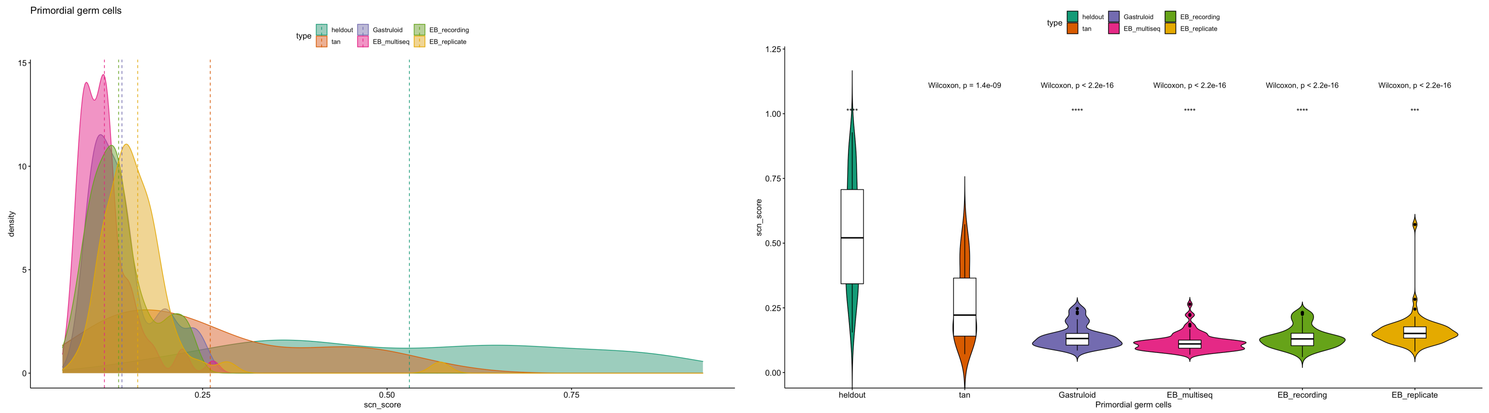


L

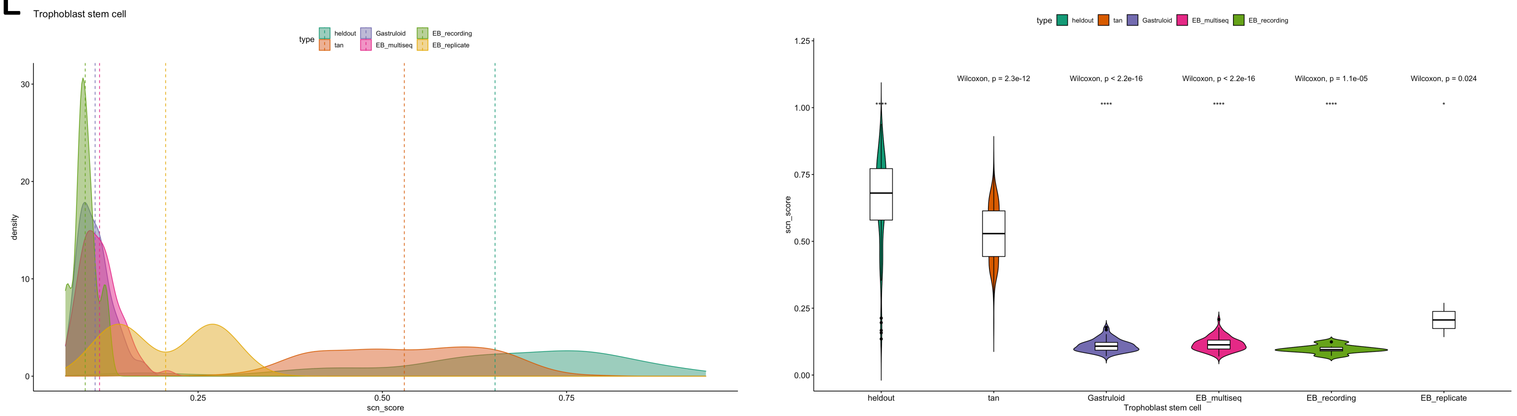

### Supplementary Figure 8

M

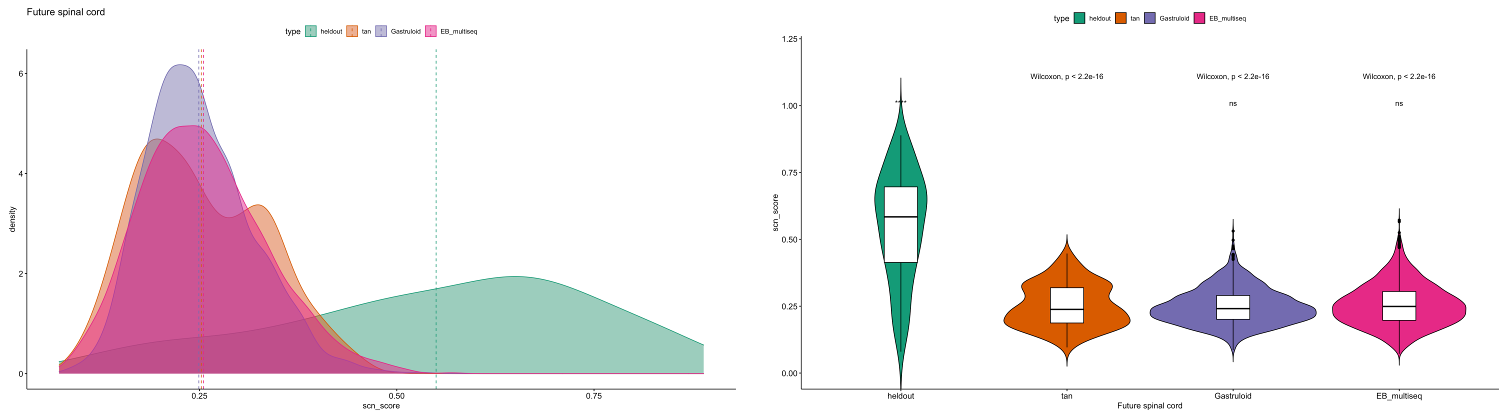


N

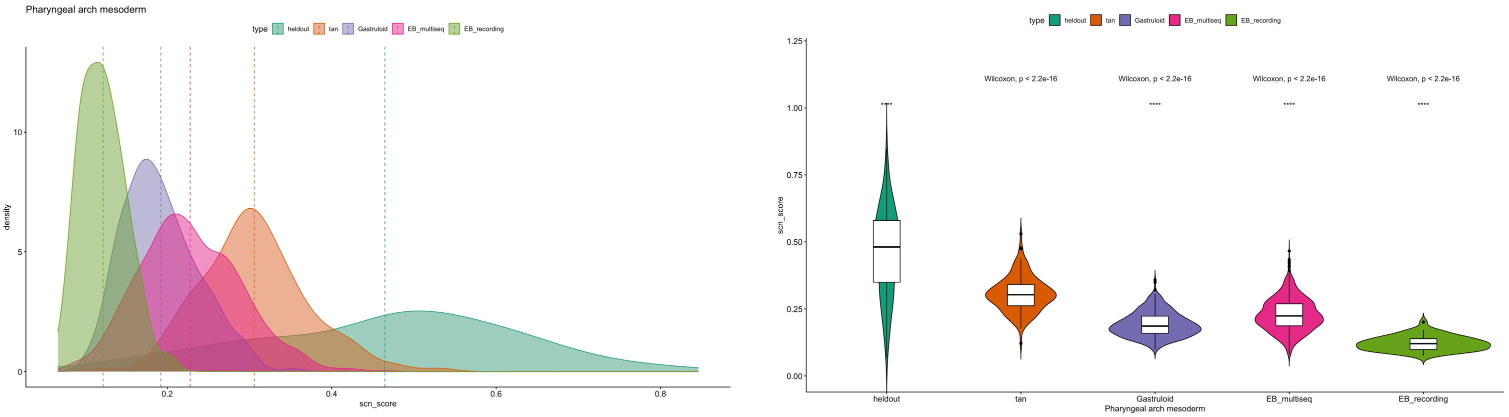


O

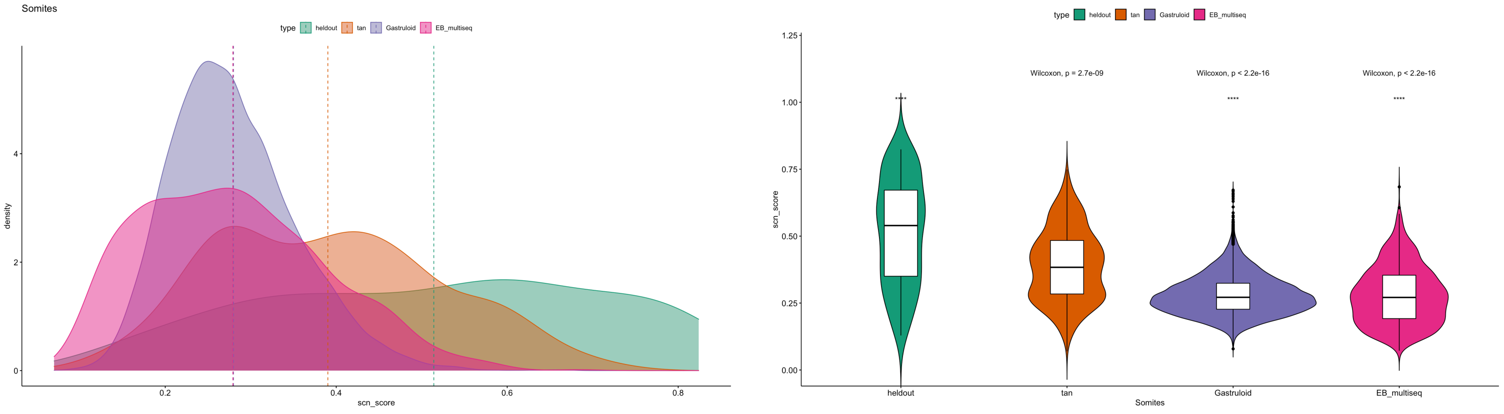

### Supplementary Figure 9

A

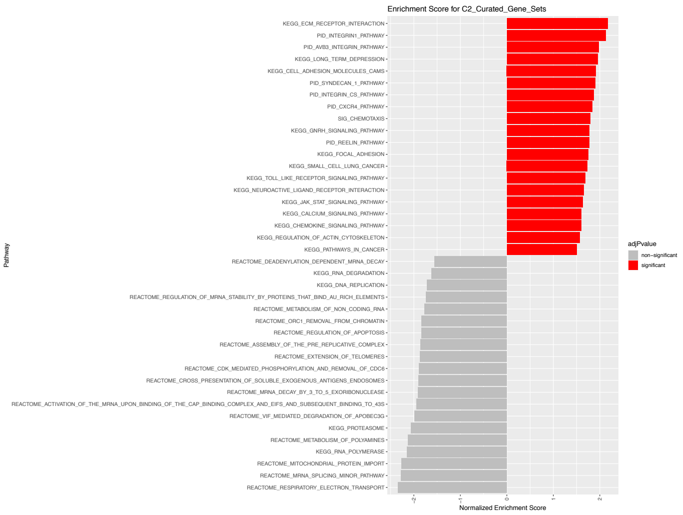


B

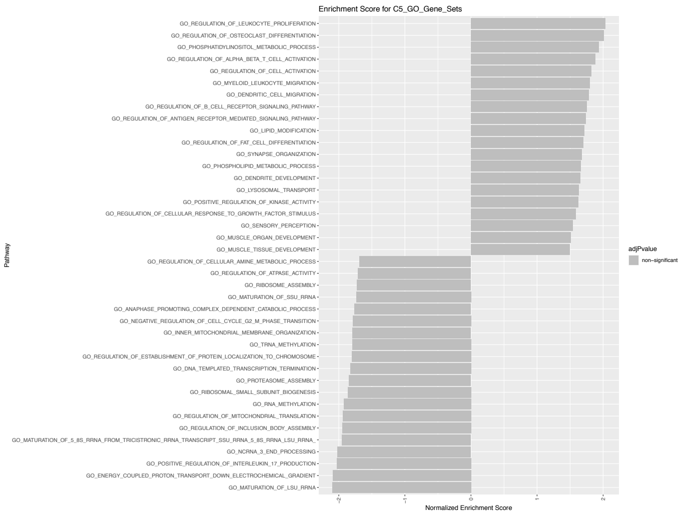


C

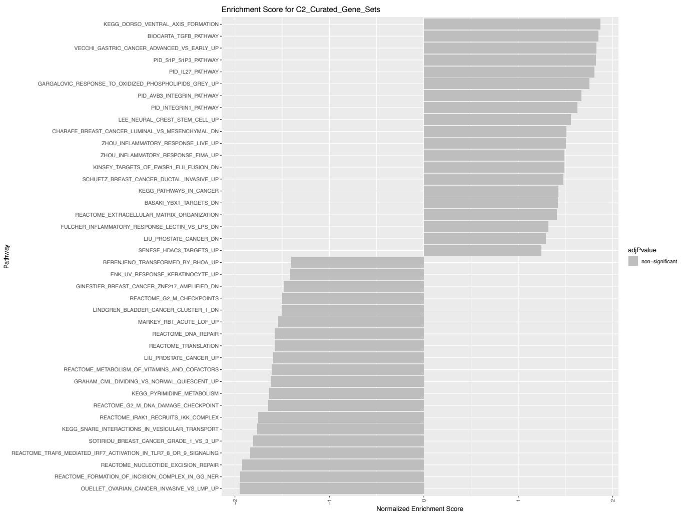


D

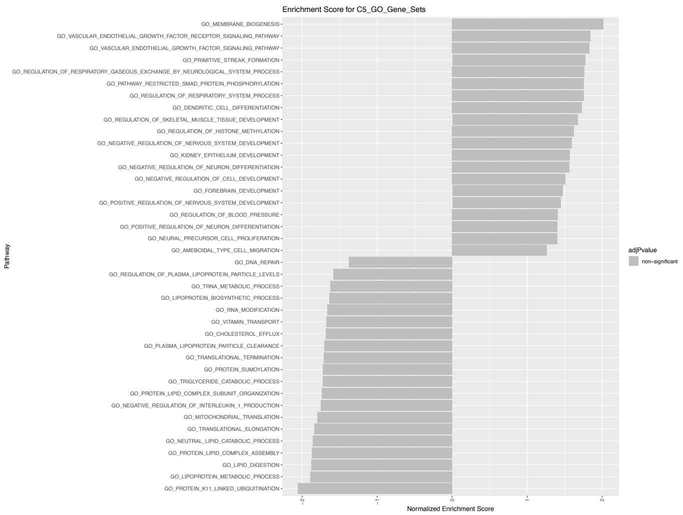

### Supplementary Figure 10

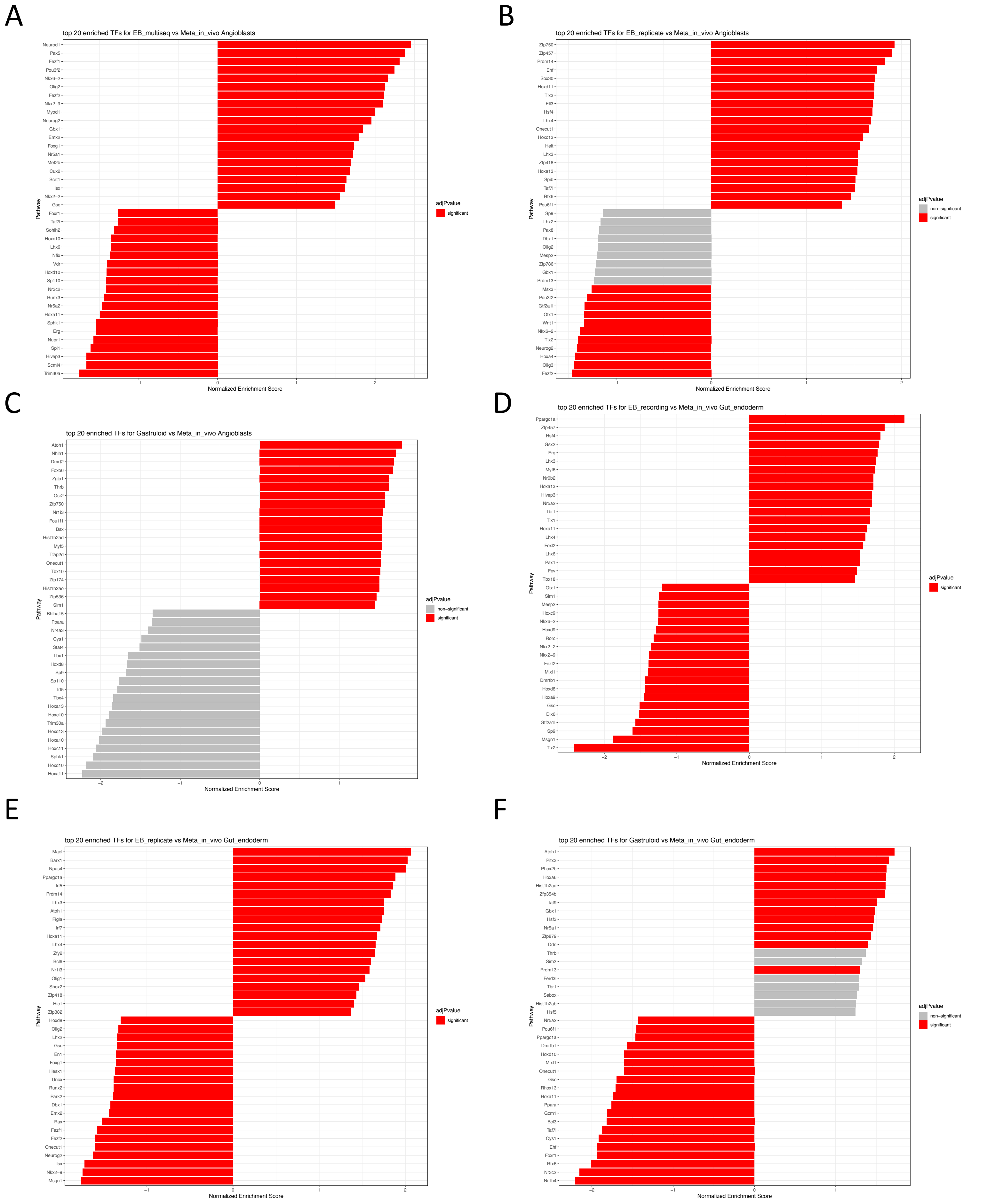
